# Extrachromosomal DNA driven oncogene spatial heterogeneity and evolution in glioblastoma

**DOI:** 10.1101/2024.10.22.619657

**Authors:** Imran Noorani, Magnus Haughey, Jens Luebeck, Andrew Rowan, Eva Grönroos, Francesco Terenzi, Ivy Tsz-Lo Wong, Jeanette Kittel, Chris Bailey, Clare Weeden, Donald Bell, Eric Joo, Vittorio Barbe, Matthew G. Jones, Emma Nye, Mary Green, Lucy Meader, Emma Jane Norton, Mark Fabian, Nnennaya Kanu, Mariam Jamal-Hanjani, Thomas Santarius, James Nicoll, Delphine Boche, Howard Y Chang, Vineet Bafna, Weini Huang, Paul S Mischel, Charles Swanton, Benjamin Werner

## Abstract

Oncogene amplification on extrachromosomal DNA (ecDNA) is strongly associated with treatment resistance and shorter survival for patients with cancer, including patients with glioblastoma. The non-chromosomal inheritance of ecDNA during cell division is a major contributor to intratumoral genetic heterogeneity. At present, the spatial dynamics of ecDNA, and the impact on tumor evolutionary trajectories, are not well understood. Here, we investigate the spatial-temporal evolution of ecDNA and its clinical impact by analyzing tumor samples from 94 treatment-naive human *IDH*-wildtype glioblastoma patients. We developed a spatial-temporal computational model of ecDNA positive tumors (‘SPECIES’) that integrates whole-genome sequencing, multi-region DNA FISH, and nascent RNAscope, to provide unique insight into the spatial dynamics of ecDNA evolution. Random segregation in combination with positive selection of ecDNAs induce large, predictable spatial patterns of cell-to-cell ecDNA copy number variation that are highly dependent on the oncogene encoded on the circular DNA. *EGFR* ecDNAs often reach high mean copy number (mean of 50 copies per tumor cell), are under strong positive selection (mean selection coefficient, *s* > 2) and do not co-amplify other oncogenes on the same ecDNA particles. In contrast, *PDGFRA* ecDNAs have lower mean copy number (mean of 15 copies per cell), are under weaker positive selection and frequently co-amplify other oncogenes on the same ecDNA. Evolutionary modeling suggests that *EGFR* ecDNAs often accumulate prior to clonal expansion. *EGFR* structural variants, including *vIII* and c-terminal deletions are under strong positive selection, are found exclusively on ecDNA, and are intermixed with wild-type *EGFR* ecDNAs. Simulations show *EGFRvIII* ecDNA likely arises after ecDNA formation in a cell with high wild-type *EGFR* copy number (> 10) before the onset of the most recent clonal expansion. This remains true even in cases of co-selection and co-amplification of multiple oncogenic ecDNA species in a subset of patients. Overall, our results suggest a potential time window in which early ecDNA detection may provide an opportunity for more effective intervention.

**Highlights:**

- ecDNA is the most common mechanism of focal oncogene amplification in *IDH*wt glioblastoma.
- *EGFR* and its variants on ecDNA are particularly potent, likely arising early in tumor development, providing a strong oncogenic stimulus to drive tumorigenesis.
- Wild-type and variant *EGFR* ecDNA heteroplasmy (co-occurrence) is common with *EGFR*vIII or c-terminal deletions being derived from *EGFR* wild-type ecDNA prior to the most recent clonal expansion.
- Tumors with ecDNA amplified *EGFR* versus *PDGFRA* exhibit different evolutionary trajectories.
- SPECIES model can infer spatial evolutionary dynamics of ecDNA in cancer.
- A delay between ecDNA accumulation and subsequent oncogenic mutation may give a therapeutic window for early intervention.

## Introduction

Genetic intra-tumor heterogeneity pervades all human cancers [1, 2]. It emerges through cell division, mutation accumulation and clonal selection, enabling somatic evolutionary processes, including mutagenesis [3] that contribute to progression and treatment resistance. Oncogene amplifications on extrachromosomal DNA (ecDNAs) are a major driver of heterogeneity [4], with random ecDNA segregation enabling extreme cell-to-cell ecDNA copy number variation upon which selection may act. ecDNA may arise early in the development of cancer, as has been shown for Barrett’s esophagus, in which the formation of ecDNA in high grade dysplasia is tightly linked to the development of esophageal cancer [5].

Glioblastoma is the most common primary intrinsic brain tumor in adults. Its prognosis has barely improved in the last few decades, with a median survival of only 14 months [6]. Targeted therapies have so far failed to make meaningful improvements to survival for glioblastoma in clinical practice [7–10]. *IDH*-wildtype glioblastomas frequently harbor oncogenes on ecDNAs, such as *EGFR*, *PDGFRA*, and *CDK4* [11–14]. It remains unclear what the evolutionary implications of these different ecDNA amplified oncogenes are in individual tumors and in tumor development. This is, in part, because random ecDNA segregation complicates unraveling evolutionary histories of ecDNA driven tumors.

Random segregation of ecDNA relative to chromosomal DNA leads to cell-to-cell copy number variability [15, 16]. The pattern of nascent RNA transcription arising from these ecDNA particles provide an opportunity to visualize the spatial organization of ecDNA across a tumor. Therefore, we developed a modeling approach called SPECIES (spatial-temporal computational model of ecDNA positive tumors) that uses structural variant analysis of focal amplifications from whole genome sequencing, DNA FISH with unbiased quantitative image analysis, RNAscope to measure nascent RNA transcription arising from oncogenes on ecDNA, and spatial-temporal computer simulations to resolve the spatial organization of ecDNA and model its evolutionary trajectories. We applied this approach to 94 *IDH*-wildtype glioblastoma samples to quantify the incidence, structure and spatial evolution of ecDNA in human glioblastoma, revealing critical underlying biological differences dictated by the oncogenes carried on ecDNA.

## Results

### The landscape of ecDNA amplified oncogenes in human glioblastomas

To determine the ecDNA landscape in human glioblastoma at clinical presentation, we first performed whole genome sequencing (WGS) and/or DNA FISH analysis of 59 adult *IDH1* wild-type, newly-diagnosed glioblastoma patient samples in the Glioblastoma-UK (GB-UK) cohort [17] (Fig. 1a; Supplementary Figure 1, see Methods for clinical details). We deciphered structural variation and ecDNA status in 49 WGS samples using AmpliconArchitect and AmpliconSuite [18, 19] (Methods). In 57% (n=28/49) of samples, we could detect at least one ecDNA amplification (Fig. 1b). Those ecDNAs contained between 2 to 52 genes with a median of 9.5 genes per ecDNA (Fig. 1i). In contrast, intrachromosomal focally amplified oncogenes were less common (n=8/49 patients, 16%), (Fig. 1b; Supplementary Figure 1). To further validate these observations, we analyzed a secondary cohort of 35 *IDH1* wild-type glioblastomas from PCAWG (Pan-cancer analysis of whole genomes, [20, 21]). Here, we detected ecDNA in 86% (n=30/35) of samples (Fig. 1b; Supplementary Figure 2), and only one case with a focal intrachromosomal amplification. All but 2 observed ecDNAs contained at least one known oncogene. In GB-UK, focal *EGFR* amplifications were always on ecDNA (n=16/49, 33%), except for one case of a breakage-fusion-bridge (BFB) cycle (copy number, CN = 6.0). Non-focal chromosomal *EGFR* amplifications were less frequent (n=5/49, 10%). Focal amplifications of other oncogenes, including *MDM2*, *CDK4*, *CDK6*, *MET*, *NMYC*, *AGAP2*, *DDIT3*, and *CLOCK* occurred exclusively on ecDNA (Fig. 1b). These results demonstrate the diversity of oncogenes that can be amplified on ecDNA in glioblastoma, and also indicate that ecDNA is overwhelmingly the primary mode of oncogene amplification in this tumor type.

**Figure 1:**
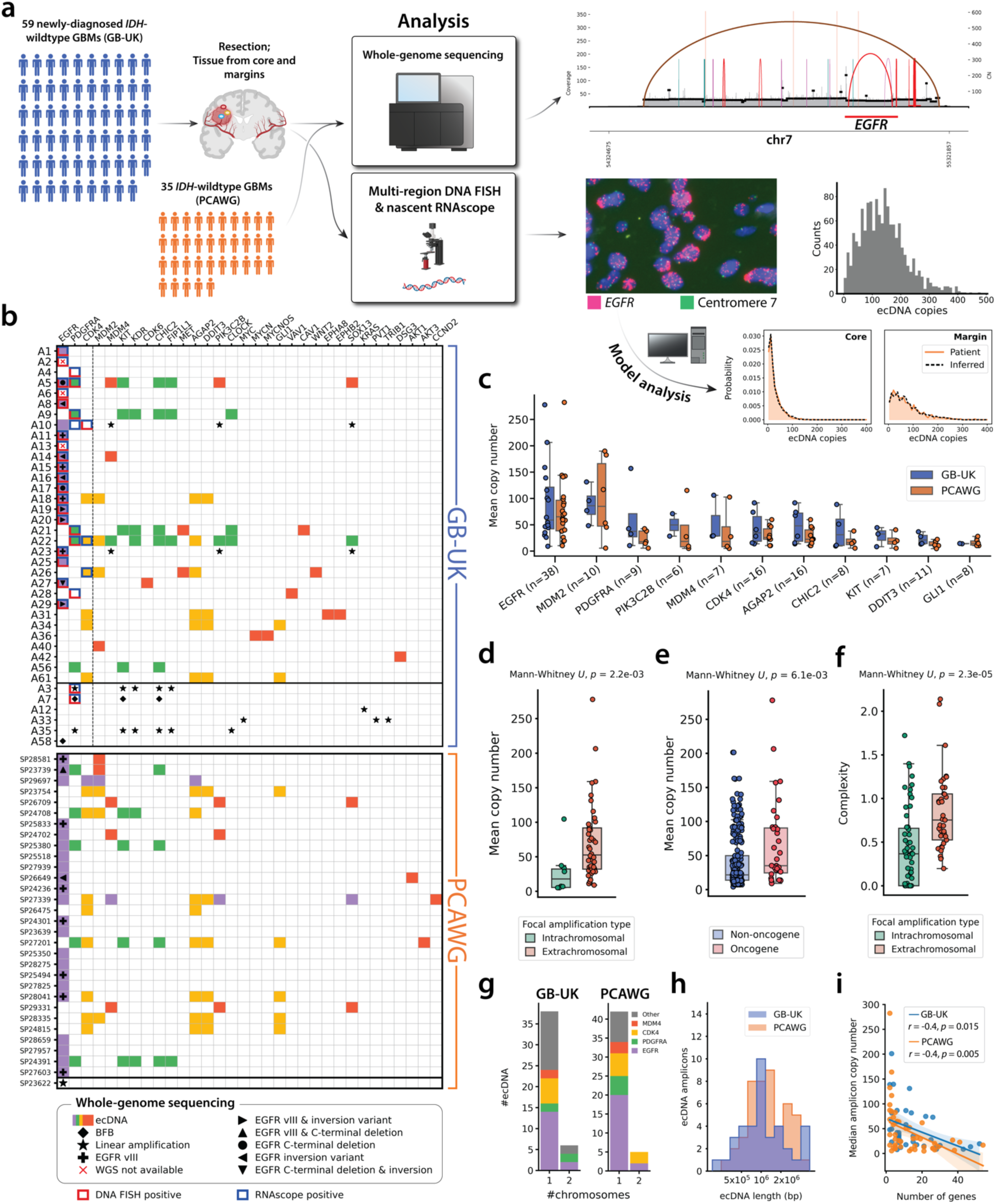
Diversity and high copy numbers of extrachromosomal DNA in IDH-wildtype glioblastomas. **(a)** Workflow of the study: 59 IDH1wt-glioblastomas (GB-UK cohort) were analyzed with DNA FISH (multi-region), mathematical modelling of single cell copy number, nascent RNAscope, and whole-genome sequencing (WGS). 35 IDH1wt-glioblastomas from the PCAWG GBM cohort were also analyzed with WGS as a validation cohort. **(b)** Waterfall plot showing focal copy number amplifications across the GB-UK and PCAWG cohorts, categorized by amplicon type (ecDNA, breakage-fusion-bridge (BFB) cycle, linear). Oncogenes with the same colored box imply they exist on the same ecDNA molecule. Horizontal line between patients A61 & A3 (GB-UK) and patients SP27603 & SP24135 (PCAWG) separate patient samples with (above) or without (below) ecDNA, detected either by whole-genome sequencing, DNA FISH or nascent RNAscope. Oncogenes tested with DNA FISH and / or nascent RNAscope, EGFR}, PDGFRA and CDK4 are separated to the left of the vertical dashed line. **(c)** Oncogenes contained within ecDNA in both GB-UK and PCAWG cohorts ranked by maximum copy number; the number of samples with each oncogene as ecDNA in the GB-UK cohort is described in brackets; only copy numbers detected by WGS are shown here. **(d)** Mean copy number of linear focal amplifications vs. ecDNA in GB-UK. **(e)** Mean copy number of non-oncogenes compared with oncogenes amplified on ecDNA in GB-UK. **(f)** Complexity score of linear focal amplifications vs. ecDNA in GB-UK. **(g)** Number of uni- and multi-chromosomal ecDNAs in GB-UK and PCAWG cohorts. **(h)** Distribution of ecDNA lengths in GB-UK and PCAWG cohorts. **(i)** Median ecDNA copy number versus number of ecDNA-amplified genes for GB-UK and PCAWG cohorts.

The glioblastoma ecDNA landscape reveals a distinctive heuristic pattern (Fig. 1b). *EGFR* containing ecDNAs do not co-amplify any other known oncogenes on the same ecDNA species. In contrast, ecDNAs containing *PDGFRA* or *MDM2* frequently co-amplify other oncogenes. In the case of *PDGFRA,* we also find *KIT*, *KDR*, *CHIC2*, *FIP1L1*, *PIK3C2B* and/or *CLOCK* on the same circular ecDNA. The native loci of these oncogenes are all on chromosome 4. Similarly, ecDNAs containing *MDM2* also harbored oncogenes *CDK4* and *AGAP2* on the same ecDNA in all cases, as well as *DDIT3* in one case, which are all genes located on chromosome 12. Consistent with previous observations [22], ecDNA amplified oncogenes can reach extremely high median copy numbers (> 100) in individual tumors (Fig. 1c) that is not seen for intrachromosomal amplifications (median ecDNA oncogene CN = 53, median intrachromosomal oncogene CN = 18; Mann-Whitney *U* test, p=0.002) (Fig. 1d). Mean copy number of ecDNA amplified oncogenes is higher compared to non-oncogenes (Mann-Whitney *U* test, p=0.006) (Fig. 1e) and ecDNAs are structurally more complex compared to intrachromosomal focal amplifications (mean complexity scores, derived using AmpliconArchitect, of 0.84 and 0.43 for ecDNA and linear amplifications respectively, Mann-Whitney *U* test, p < 0.0001) (Fig. 1f). Most ecDNAs contain DNA from a single chromosome. We find ecDNA derived from two chromosomes (translocation-ecDNAs) in 12% (n=6/49) of GB-UK and 14% (n=5/35) of PCAWG samples (Fig. 1g). We did not detect combinations of 3 or more chromosomes on the same ecDNA.

Interestingly, ecDNA sizes are conserved within one order of magnitude both in GB-UK and PCAWG (minimum = 180 kbp; maximum = 5.52 Mbp) (Fig. 1h). We furthermore observe a negative correlation between the number of ecDNA-amplified genes and ecDNA copy number (Pearson *r* = −0.4; *p* = 0.015 and *r* = −0.4; *p* = 0.005 for GB-UK and PCAWG respectively) (Fig. 1i), suggesting the possibility that ecDNAs evolve over time. This is further supported by ecDNA structure. For example, all ecDNAs in GB-UK containing *CDK4* and *MDM2* lost the intervening chromosomal segment between both genes, suggesting balancing selection to minimize ecDNA size while conserving oncogenes.

### Spatial ecDNA copy number heterogeneity

Although *SEC61G*, a gene promoting immune evasion [23], co-amplifies on the same ecDNA with *EGFR*, no other known oncogenes are in the proximity of *EGFR* that could be co-amplified on ecDNA partially explaining the observed patterns of different ecDNA amplified oncogenes. However, the frequent convergent evolution of extremely high ecDNA amplified *EGFR* in glioblastoma suggests an important role in tumor evolution and that *EGFR* and its variants are under strong selective pressure. If this were true, we would anticipate different patterns of spatial temporal evolution of *EGFR*-containing ecDNAs, relative to ecDNAs that contain other oncogenes. To test this hypothesis, we measured the spatial variation of the two most common ecDNA amplified oncogenes, *EGFR* and *PDGFRA,* with DNA FISH in the core and infiltrating margin in each GBM in the GB-UK cohort (Fig. 2a, b). The DNA FISH and WGS derived mean ecDNA copy numbers strongly correlate (Pearson r = 0.6, p = 0.01), (Supplementary Figure 3). Unbiased image analysis (Methods), then allowed us to quantify the per-cell ecDNA count for each core and margin region separately. Both *EGFR-* and *PDGFRA*-ecDNAs, when observed in tumor cores, are also frequently seen in margins and leading edges (72%, n=23/32; Supplementary Figure 4). ecDNA copy numbers show the typical wide cell-to-cell variation that emerges from random ecDNA segregation during cell division[15, 16]. However, *EGFR*-ecDNA single-cell copy number distributions are highly variable across different regions of a patient’s tumor. Typically, distributions in core samples are wider and shifted towards higher mean copy number compared to margin samples of the same tumor. In contrast, *PDGFRA*-ecDNA copy number distributions are more similar between core and margin samples of the same tumor (Fig. 2a, b).

**Figure 2:**
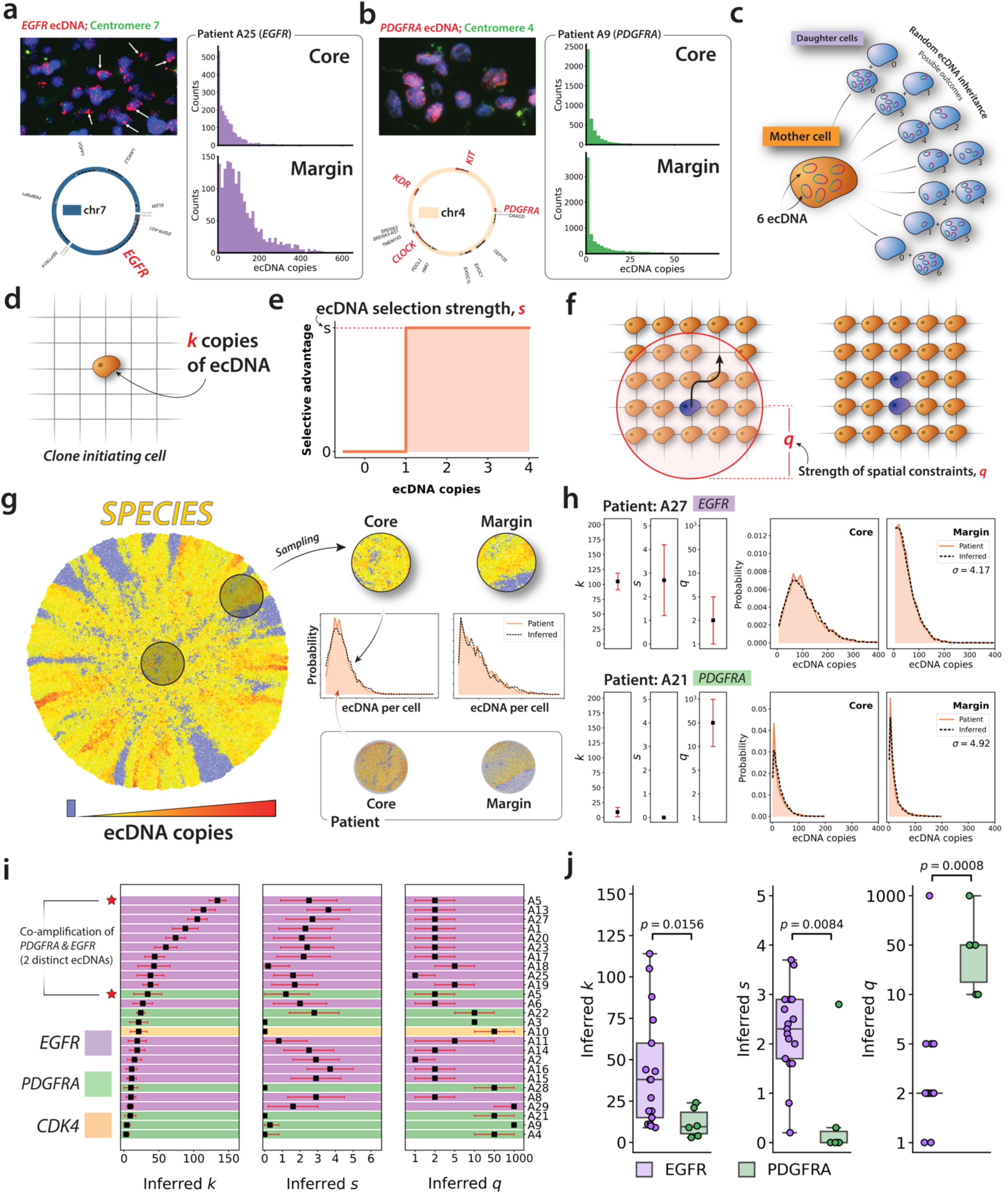
SPECIES Spatial modelling of ecDNA evolution. **(a-b)** DNA-FISH staining of two representative GBM samples from GB-UK cohort, revealing **(a)** *EGFR*- and **(b)** *PDGFRA*-ecDNA. *EGFR*- ecDNA appear congregate into “hubs” (white arrows). Single-cell ecDNA copy number distributions are derived from DNA FISH images using unbiased image analysis. **(c)** During cell division, following replication of ecDNA, amplicons are divided binomially amongst daughter cells. **(d)** SPECIES is initiated with a single tumor cell, containing n copies of ecDNA. **(e)** Model of ecDNA selection. Cells carrying 1 or more copies of ecDNA divide at a rate *s* faster than cells lacking ecDNA **(f)** Cells push neighbors within a radius *q* on the lattice to make space necessary for cell division. **(g)** Spatial computational model of ecDNA-driven tumors. Approximate Bayesian computation (ABC) was used to infer optimal model parameters for initial number of ecDNA, *k*, ecDNA-conferred selection, *s*, and cell pushing strength, *q*, for a given set of patient tumor measurements. **(h)** Example outputs of ABC model fitting algorithm, shown for patients A27 and A21. Inferred *s*, *k* & *q* (top row) and model best-fit distributions from simulated tumors (bottom row). Sum of Wasserstein distance between patient and simulated distributions for tumor core and infiltrating margin, representing closeness of fit, is denoted by *σ*. **(i)** Summary of inferred model parameters for all sampled human GBM tumors. Highlighted color represents ecDNA amplified oncogene (*EGFR*, *PDGFRA* or *CDK4*) in each tumor. **(j)** Inferred model parameters for all patients, stratified by ecDNA-amplified oncogene (*EGFR* or *PDGFRA*).

We then tested how such differences emerge in the spatial-temporal evolution of ecDNA positive tumors. We developed stochastic computer simulations of expanding ecDNA driven tumor populations (SPECIES, *SP*atial *EC*dna Intratumor Evolution Simulation), using a lattice-based cellular automaton model (Methods) (Fig. 2c-f) [24, 25]. SPECIES requires three essential parameters, the ecDNA copy number *k* in the first tumor initiating cell, the strength of the ecDNA- conferred replicative advantage, *s,* and the range of spatial constraints, *q* (Fig. 2d-f). For any set of these parameters, SPECIES generates explicit spatial and temporal patterns of single cell ecDNA copy numbers, enabling us to mimic *in vivo* multi-region sampling (Fig. 2g). Different combinations of the parameters *k, s* and *q* result in dramatically different spatial patterns of ecDNA cell to cell variation (Supplementary Figure 5). Large *q* values, leading to weak spatial constraints, produce more spatially homogeneous tumors, whereas small *q* gives rise to local spatial sectors and greater spatial variation in ecDNA copy number. Despite our chosen model of ecDNA- conferred selection favoring equally high and low ecDNA copy number cells (constant selection model, see Methods), larger *s* nevertheless drives the tumor population to higher ecDNA copy numbers, through ecDNA-positive cells outcompeting ecDNA-free cells. Simulated spatial ecDNA patterns are most sensitive to changes in selection strength, *s,* when combined with low *k*, since tumors initiated with few ecDNA copies (i.e. low *k*) rapidly generate less fit ecDNA-free cell lineages, as a result of random ecDNA segregation. However, due to the inherent stochasticity of cell divisions and random ecDNA segregation, individual realizations of simulations vary in the exact spatial patterns for any fixed set of these evolutionary parameters.

To quantitatively estimate which parameter combinations describe tumor growth dynamics best in individual patients, we implemented approximate Bayesian computation (ABC) [26, 27] to fit SPECIES simulations to patient-derived ecDNA copy number distributions (Methods) (Fig. 2h; Supplementary Figures 6-12). SPECIES accurately recapitulates ecDNA copy number distributions in core and margin samples for each patient (Wasserstein distance metric *σ* < 10, see Methods). The best inferred model parameters differ between patients. We find a wide range of ecDNA copy numbers *k* in the first tumor initiating cell, ranging from 2 (patient A4, *PDGFRA*) to 134 (patient A5, *EGFR*). In 15/26 tumors, ecDNA copy number distributions suggest a model *k* value of 20 or more. This implies that, in these patients, ecDNA are likely to have been present some time prior to the most recent tumor clonal expansions. Evidence for the accumulation of ecDNA in pre-malignant populations recently emerged in longitudinal studies of cancerous transformations in Barrett’s esophagus [5]. In 17/26 tumors we estimate very strong selective advantages for ecDNA+ cells (*s* ≥ 1.6) (16/18 *EGFR*; 1/7 *PDGFRA*; 0/1 *CDK4*). 9 samples are consistent with either weak positive or neutral dynamics of ecDNA positive cells (error regions on inferred *s* values for these patients include *s =* 0), though the high abundance of ecDNA observed in these 9 glioblastomas further suggests positive ecDNA-conferred fitness advantage [16]. In most cases, SPECIES inferences suggest strong spatial constraints, corresponding to small *q* (*q* = 2 and *q* = 5), implying that tumor cells are constrained for space during tumor expansion, but are not strictly confined to boundary growth. In summary, evolutionary modeling suggests the majority of oncogene-containing ecDNAs to be important cancer drivers that must have been present at the onset of the most recent clonal expansion.

### EcDNA oncogene-specific glioblastoma evolution

We then asked whether glioblastoma evolutionary dynamics depend on ecDNA cargo. In GB-UK, DNA FISH derived copy numbers of *EGFR-* versus *PDGFRA*-ecDNA had significantly different distributions for both maximum and mean copy number, as well as cell-to-cell variance and skew (Supplementary Figure 13, Mann-Whitney *U* test, *p* = 6.4 × 10^-4^, *p* = 2 × 10^-5^, *p* = 2.7 × 10^-4^ and *p* = 4.8 × 10^-4^ respectively). In keeping with these differences, SPECIES predicts a wider range of inferred *k* and *s* parameters amongst tumors with *EGFR*-ecDNA compared to *PDGFRA-* ecDNA (interquartile range of *k* was 45 and 13 for *EGFR* and *PDGFRA* respectively; interquartile range of *s* was 1.0 and 0.2 for *EGFR* and *PDGFRA* respectively, Fig 2i). We observe different evolutionary dynamics between *EGFR*- and *PDGFRA-*ecDNA amplifications. Tumors containing ecDNA-amplified *EGFR* show significantly larger predicted initial ecDNA copy number *k*, selective strength *s*, and stronger spatial constraints, *i.e.* smaller *q* (Mann-Whitney *U* test, *p* = 0.0156, *p* = 0.0084 & *p* = 0.0008 respectively) compared to tumors with ecDNA-amplified *PDGFRA* (Fig. 2j). These predicted oncogene-level differences are independent of variations in the spatial location of the core sample (to reflect similar variances from the patient tissue acquisition), and different cell birth/death ratios (Supplementary Methods; Supplementary Figure 14). Furthermore, model fit accuracy declines noticeably when restricting initial simulated ecDNA copy numbers to *k* = 1. This decline is most pronounced in patient samples with the highest predicted *k* values, further suggesting that in some patients a high initial ecDNA copy number *k* is required to capture observed shifts of ecDNA distributions between core and margin samples (Supplementary Figure 15).

### Evolution of *EGFR* ecDNA variant heteroplasmy

What then determines the transition of pre-malignant *EGFR* ecDNA accumulation into clonal expansions? A two-hit model of ecDNA formation followed by a secondary event initiating clonal expansion could potentially explain high initial *EGFR* ecDNA copy numbers. To see if there is evidence for a second hit, we analyzed the structural variants (SVs) on *EGFR* ecDNA and its variation within single tumors (Supplementary Figure 16). In GB-UK 75% (*n*=12/16) of patients with ecDNA amplified *EGFR* carried mutated *EGFR* variants on ecDNA, with *n*=7/16 harboring the GBM specific oncogenic *EGFRvIII* variant (constitutively active *EGFR* [28], Fig. 3a). *EGFRvIII* was exclusively amplified on ecDNA. Surprisingly, in all patients *EGFRvIII* co-existed with *EGFR* wild-type (*EGFR*wt) ecDNA, usually at lower copy numbers compared to *EGFR*wt amplifications. *EGFR*wt and *EGFRvIII* share the same DNA breakpoints suggesting *EGFRvIII* ecDNA variants may arise from *EGFR*wt ecDNAs. A similar dynamic has been observed previously in the GBM39 cell line using CRISPR-CATCH [29]. In addition, three glioblastomas carried ecDNAs with an *EGFR* c-terminal deletion (exons 25-27), an activating *EGFR* truncation ([30]; [31]) not previously reported to occur on ecDNA. Four glioblastomas had an inversion SV between exons 1 and 7, that codes for the *EGFR* extracellular binding domain and is likely to lead to a non-transcribed segment with functional equivalence to *EGFRvIII* (Supplementary Figure 17). This heteroplasmy-like variation of *EGFR* structural variants (*i.e.* their co-occurrence with *EGFR*wt) is enabled by the wide cell to cell ecDNA copy number heterogeneity and must emerge from the interplay of random segregation, selection and mutation of ecDNAs. As the ecDNA is characterized in a single head-to-tail structural variant, the most likely scenario for *EGFRvIII* emergence is first through generation of *EGFR*wt ecDNA, which amplifies before subsequently acquiring a *EGFRvIII* variant. It remains unclear, however, if the mutating event occurs before or after the initial expansion of the tumor.

**Figure 3:**
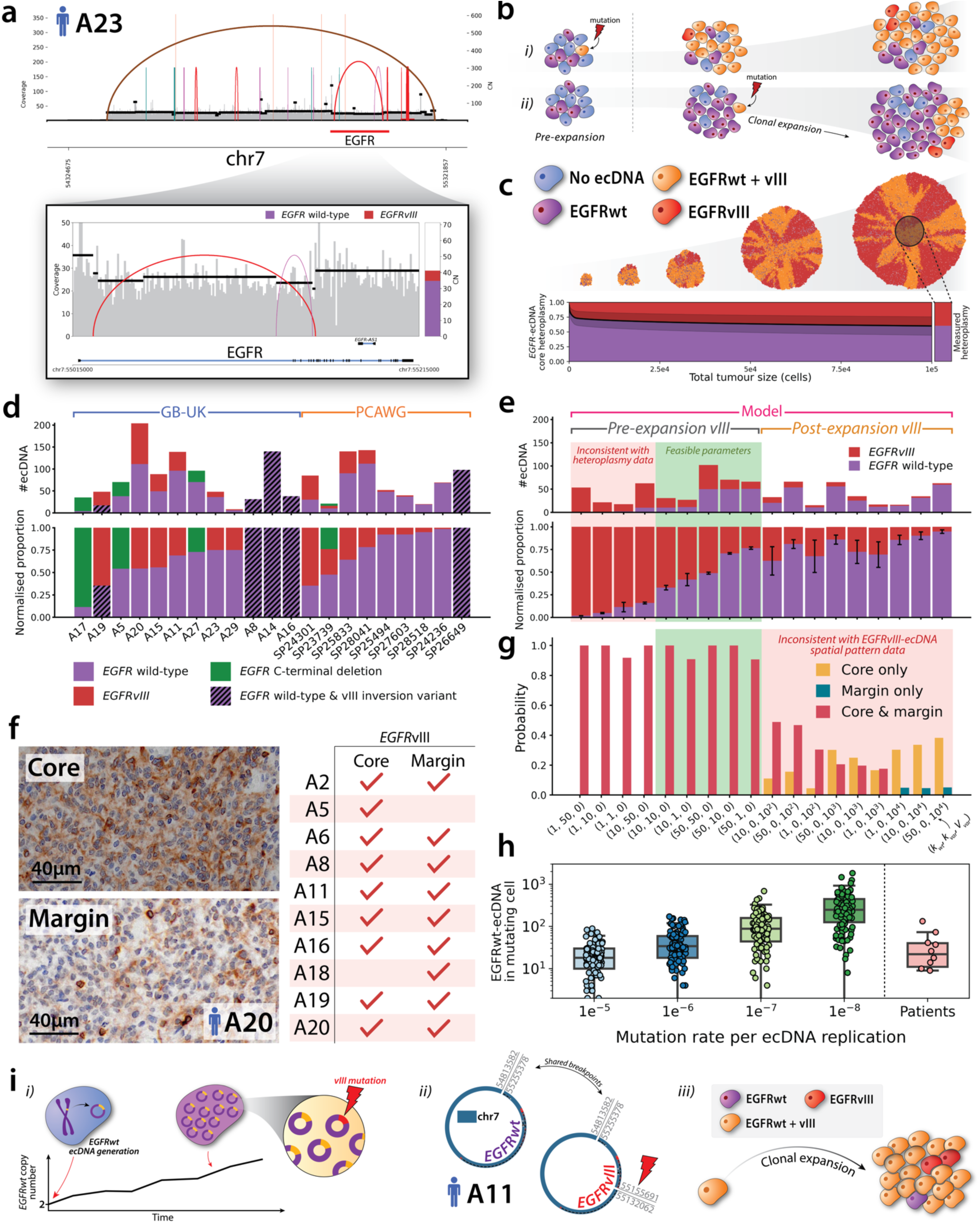
Selection dynamics of *EGFR* structural variants on ecDNA. **(a)** A subclonal *EGFRvIII* variant on ecDNA in patient A23. **(b)** Effects of (i) pre- and (ii) post-clonal expansion mutation of *EGFR*-ecDNA on its resulting spatial pattern in the expanded tumor. **(c)** SPECIES was applied to simulate the emergence of an advantageous *EGFRvIII* mutation on ecDNA, either prior to or after the onset of clonal expansion of the tumor. Example shown is for a post-expansion mutation, with parameters (*k*_*wt*_ = 20, *k*_*var*_ = 1, *V*_*var*_ = 0). Standard deviation of ecDNA heteroplasmy shown by shaded region in lower plot. **(d)** Variant heteroplasmy in the core region of all samples across GB-UK and PCAWG cohorts which contained EGFR variant-bearing ecDNA. **(e)** Model predictions for resulting variant heteroplasmy in tumor for a range of wild-type and variant bearing ecDNA copy numbers in the clone-initiating tumor cell (for pre-expansion EGFR mutation) and mutation times (for post-expansion EGFR mutation). Red shaded region denotes parameter combinations inconsistent with observed patient core heteroplasmy values. Mean +- variance derived from 1000 simulated tumors. **(f)** (*Left*) Immunohistochemistry (IHC) slide for patient A20, showing presence of *EGFRvIII* in the tumor core and infiltrating margin (blue = haematoxylin; brown = *EGFRvIII*). (*Right*) Summary of *EGFRvIII* in core and margins of GB-UK cohort samples, determined by IHC. **(g)** SPECIES model predictions for the probability of observing *EGFRvIII* ecDNA across tumor core and margin, for a pair of core and random margin regions. Mean values derived from 1000 simulated tumors. **(h)** Distributions of EGFRwt-ecDNA abundance in the cell which acquires the first mutation, for a range of ecDNA mutation rates. Patient values derived from their respective inferred *k* values, by assuming that the arrival of a mutation on ecDNA initiates clonal expansion, implying that the number of EGFRwt-ecDNA in the mutating cell is thus equal to *k*-1 (with the remaining 1 ecDNA carrying the mutated gene). **(i)** Timeline of EGFR- ecDNA tumorigenesis. (i) Initial EGFRwt-ecDNA is generated and accumulates in pre-malignant cells, conferring a moderately positive selective advantage. In time, EGFRvIII mutation occurs on one of the accumulated EGFRwt-ecDNAs, conferring a strong positive advantage. (ii) Reconstructed ecDNA structures from GB-UK patient A5 confirms EGFRwt and EGFRvIII ecDNAs share common breakpoints, indicating EGFRvIII arose from mutation of an existing EGFRwt-ecDNA. (iii) EGFRvIII-ecDNA further promotes clonal expansion, driving tumor growth.

To better understand the dynamics of activating *EGFR* mutations on ecDNA, we adapted SPECIES to simulate expansions of advantageous ecDNA variants in the background of pre-existing *EGFR*wt ecDNA (Fig. 3b, c). This model allows us to study how the initial copy numbers of wild-type and mutated *EGFR*-ecDNA, *k*_*wt*_ and *k*_*var*_ respectively, and the time of mutant ecDNA emergence, *V*_*var*_, affect ecDNA heteroplasmy (defined here as the percentage of ecDNA copies which are amplifying *EGFR*wt, see Methods). Simulated and patient *EGFR*wt and *EGFRvIII* ecDNA copy numbers are consistent with the emergence of an activating variant (*vIII* or c-terminal deletion) at or close to the onset of clonal expansion (Fig 3d, e). Simulations with either a large initial abundance of *EGFR*wt (*k*_*wt*_:*k*_*var*_ ratio of 10:1, 50:1 or 50:10), or both ecDNA types (*k*_*wt*_:*k*_*var*_ ratio of 10:10 or 50:50), or where mutant *EGFR*-ecDNA was not present at the onset of clonal expansion but occurred in its early stages (*V*_*var*_ ≤ 10^4^ cells), are compatible with patient data. Simulations of tumors derived from a cell with low *EGFR*wt copy number (*k*_*wt*_:*k*_*var*_ ratio of 1:1, 1:10, 1:50 or 10:50, Fig. 3e, red shaded region) did not match qualitatively with observed copy numbers in patient samples.

To further narrow down the timing of *EGFR*-ecDNA variants, we measured the spatial extent of the *EGFRvIII* ecDNAs in the GB-UK cohort using immunohistochemistry (IHC) (Methods) (Fig. 3f). *EGFRvIII* ecDNAs are typically maintained across the core and infiltrating margin of tumors (n=8/10) which, according to spatial simulations, strongly suggests that *EGFRvIII* ecDNA variants emerge prior to tumor clonal expansion and are present in the tumors most recent common ancestor cell (Fig. 3g). SPECIES simulations with later arising *EGFRvIII* ecDNAs give rise to spatially variegated patterns of *EGFR*wt and *vIII* ecDNA, with large regions of the tumor margin only harboring one but not both types (Fig. 3g, red shaded region).

Spatial simulations of *EGFR* heteroplasmy evolution and ABC parameter inferences together suggest the following typical scenario of ecDNA amplified *EGFR* in human GBM: *EGFR*wt-ecDNA forms in pre-malignant tissue, conferring a moderate selective advantage (Fig. 3i (i)). In time, *EGFR*wt-ecDNA copy numbers increase, further increasing the risk of acquiring a strongly advantageous activating mutation on *EGFR*wt-ecDNA (*vIII* or a c-terminal deletion). Shared breakpoints across the *EGFRwt* and *vIII* ecDNA demonstrate categorically that the *vIII* variant occurred on one of the *EGFRwt*-ecDNA accumulated during this period (Fig. 3i (ii)). Simulations of *EGFR*wt-ecDNA dynamics in pre-malignant tissue confirm that an *EGFR* activating mutation on ecDNA typically arises in a cell with large abundance (>10 copies) of *EGFR*wt-ecDNA, giving rise to an initial heteroplasmy of 90% or above (Fig. 3h). During clonal expansion, *EGFR*wt-ecDNAs hitchhike on strongly selected *EGFRvIII-*ecDNA (*i.e.* benefit from the evolutionary advantage of the more favorable *EGFRvIII*-ecDNA by virtue of their co-occurrence within cells), maintaining intra-tumor *EGFR* heteroplasmy (Fig. 3i (iii)). This would be consistent with previous reports of cooperativity between *EGFRwt* and *EGFRvIII* in malignant transformation [32]. Moreover, multiple SVs consistent with *vIII* are observed in samples A20 from GB-UK, and SP24301, SP23739 and SP28041 from PCAWG, suggesting ongoing evolution of activating *EGFR* mutations within these ecDNAs (Supplementary Figure 16).

This causal sequence is consistent with all our patient samples except two. Patient A5 harbored *EGFRvIII*-ecDNA only in the tumor core, and so a *EGFR*wt-ecDNA may have mutated not before, but shortly after onset of clonal expansion (Fig. 3g). Similarly, patient A18 harbored *EGFRvIII*- ecDNA only in the infiltrating margin, perhaps suggesting the *vIII* mutation developed on ecDNA late into clonal expansion, at the expanding edge of the tumor, such that only *EGFR*wt-ecDNA were present in the tumor core.

### Co-selection and co-segregation dynamics of multiple ecDNA species

Recently, the distinct nature of co-amplification and co-selection of multiple ecDNA species in cancers has been discovered [29]. Here, 16% (*n=*8/49) of GB-UK GBMs harbored at least two independent oncogenes amplified on different ecDNA. In one case (patient A5), the tumor contained 3 distinct ecDNA species which together amplified 7 oncogenes, including *EGFR*, *PDGFRA* and *MDM4* (Fig. 4a, b). The tumor of patient A21 harbored one ecDNA containing *PDGFRA / KIT / KDR*, and a second ecDNA with *MET* (CN = 11 and 13 respectively). Amongst the GB-UK GBMs with multiple ecDNA species, n=5/8 contained *MDM2* or *MDM4* amplified on one of the ecDNAs, implying that loss of *TP53* pathway may facilitate ecDNA formation with multiple ecDNA species (Fig. 4c). Similarly, in PCAWG, we observed multiple ecDNAs per tumor in 31% (n=11/35) of cases, with combinations such as *EGFR* ecDNA with a *CDK4* / *MDM2* ecDNA. The data suggesting that disruption of TP53 may be an important event in facilitating ecDNA formation, is consistent with the recent observation in patients with Barrett’s esophagus who progressed to esophageal cancer, for which *TP53* loss always preceded ecDNA development [5]. In patients with multiple ecDNA oncogenic species, their nascent expression was frequently seen in different cells in a single tumor (Methods), suggesting the presence of more than one ecDNA species can further increase intercellular oncogene expression heterogeneity.

**Figure 4:**
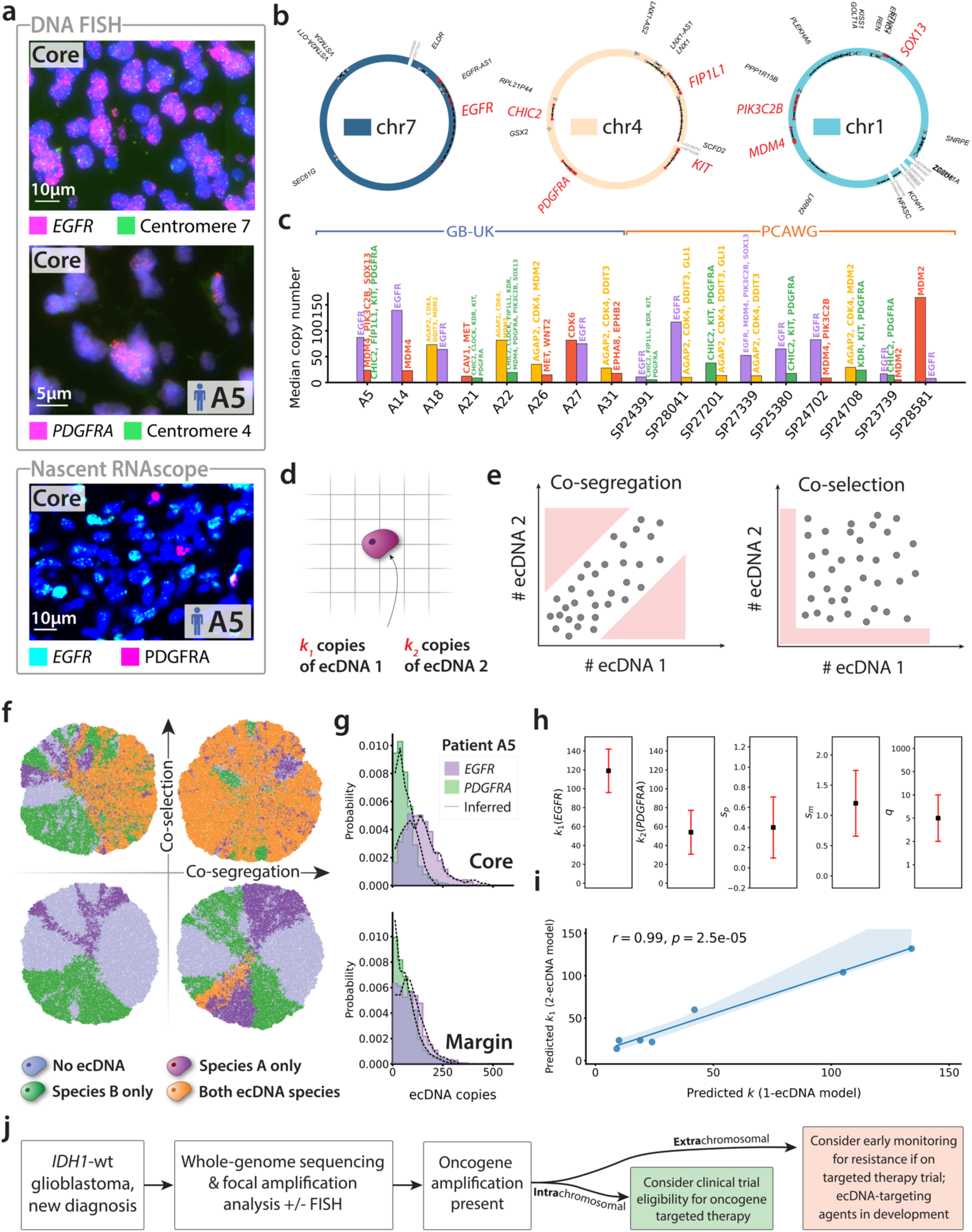
Simulating multi-ecDNA dynamics *in vivo.* **(a)** DNA-FISH and nascent RNAscope images from patient A5, showing coexistence of both *EGFR*-ecDNA and *PDGFRA*-ecDNA. **(b)** Circular structures of 3 distinct oncogenic ecDNAs detected in patient A5. **(c)** Glioblastomas in GB-UK and PCAWG cohorts containing more than one oncogenic species and their respective copy numbers. **(d)** In a multiple-ecDNA application of SPECIES, the clone-initiating cell begins with *k*_1_ and *k*_2_copies of ecDNA 1 and ecDNA 2 respectively. **(e)** Representation of the impacts of ecDNA co-segregation and co-selection on number of inherited ecDNA copies during cell division. Co-segregation drives correlated inheritance of both ecDNA types, whilst co-selection favors tumor cells carrying at least one copy of each type. **(f)** Application of SPECIES to simulate two separate ecDNA species. Simulated tumors are initialized with a single cell, carrying *k*_1_ and *k*_2_ copies of each ecDNA species. Representative model images showing examples of low and high co-segregation and co-selection. **(g-h)** Parameter inference summary for patient A5, for which we measured both the *EGFR*-ecDNA and *PDGFRA*-ecDNA copy number distributions using DNA-FISH, using multiple-ecDNA SPECIES. Parameters *s*_*p*_ and *s*_*m*_ represent selection coefficients for tumor cells with 1 (pure) or 2 (mixed) ecDNA species respectively. **(i)** Comparison of inferred *k* and *k*_1_ values for all patients confirmed by whole-genome sequencing to harbour two or more ecDNA species. **(j)** The presence of ecDNA amplifications may be used to aid stratification, given oncogenes on ecDNA have an inherent resistance mechanism through the ability of ecDNA to dynamically adjust copy number in response to targeted agents. Earlier monitoring and intervention is recommended for those patients with ecDNA that will have targeted therapies.

SPECIES provides an opportunity to probe the dynamics of multiple ecDNA species in tumors. In patient A5, DNA FISH revealed two separate ecDNA amplicons, amplifying *EGFR* and *PDGFRA* (Fig 4a, b). When inferring tumor dynamics in this patient, we did so both for the *EGFR* and *PDGFRA* separately, thus providing two sets of *k*, *s*, and *q* values. As expected, best-fit values for *q* using data for either ecDNA amplicon in this patient agree, given this parameter models the growth in the tumor and does not describe ecDNA specific dynamics. Inferences on the *EGFR* ecDNA copy number distribution resulted in the largest predicted value of *k* (*k* = 134 ± 12) across the entire cohort. Furthermore, for this patient, SPECIES predicted the largest value of *k* and the lowest value of *q* across all *PDGFRA*-amplifying tumors (*k* = 34 ± 20; *q* = 2), suggesting that the overall dynamics may be dominated by the *EGFR* amplified ecDNA.

However, the basic framework of SPECIES discussed above does not account for emergent dynamics such as co-segregation (correlated inheritance of both ecDNA species) and co-selection (selective advantage for maintaining a mixture of both ecDNA species within a cell) of distinct ecDNA species. These mechanisms may provide an alternative explanation to the inferred high abundance of pre-existing ecDNAs in some tumors. To model the dynamics of multiple ecDNA species more faithfully, we adapted SPECIES to include two distinct ecDNA types and modeled co-segregation and co-selection using the approach of Hung *et al*. [29](Supplementary Methods) (Fig. 4d, e). Depending on the strength of co-segregation and co-selection of two ecDNAs, different scenarios for ecDNA maintenance across tumor space emerged (Fig 4f). DNA FISH and nascent RNAscope analysis revealed that GB-UK GBMs harboring multiple ecDNA species, A5, A10 & A22, each maintained these populations in tandem across the core, infiltrating margin and leading edge of the tumor, except for the infiltrating margin of A10 (Supplementary Figure 4). These observations are consistent with coordinated inheritance and co-selection of both ecDNA types (Fig. 4f). To further explore this scenario, we fit the multiple-ecDNA SPECIES model to the multiple ecDNA GBMs in GB-UK for which we had single-cell ecDNA copy number distributions (n=7/8 patients). In each case, we compared the inferred initial copy number for one of the amplicons to that inferred with the single-ecDNA SPECIES model (Supplementary Methods). The inferred initial ecDNA numbers of *EGFR*- and *PDGFRA*-ecDNA in patient A5 were highly consistent across the single-versus multiple-ecDNA SPECIES models (Fig. 4g, h; Supplementary Figure 18). Across all co-amplified patient tumors in GB-UK, the initial ecDNA copy number inferred with both SPECIES models were strongly correlated (Fig. 4i, Pearson *r* = 0.99; *p* = 2.5 × 10^!#^). Thus, while co-segregation and co-selection dynamics likely play a role in these tumors to maintain high fractions of mixed cells, a high starting ecDNA copy number for one of the ecDNA species is still required to explain the observed spatial patterns of the ecDNA copy number distribution.

## Discussion

In summary, our paper reveals four critical insights about the role of ecDNA in *IDH*-wt glioblastoma. First, focal oncogene amplification almost always occurs on ecDNA. Copy number distributions of ecDNA across spatially distinct regions vary between patients and ecDNA cargo, suggesting that different oncogenic ecDNAs are under varying degrees of positive selection with spatial growth constraints and oncogene-specific ecDNA dynamics.

Second, it shows that *EGFR* and its variants are particularly potent, likely arising early in tumor development, and likely providing a sufficient oncogenic stimulus to drive tumor formation and progression. Previous molecular analyses and mouse models have implicated *EGFR* amplification as an early or transformational event in some glioblastomas [33–36], though without consideration of ecDNA; here, we advance this work to quantitatively account for ecDNA spatial dynamics, revealing two groups of EGFR tumors - those with pre-malignant accumulation of ecDNA where ecDNA is likely to contribute to initiating clonal expansion, and those tumors with few ecDNAs in early tumorigenesis. The emergence of high-level *EGFR* amplification facilitates subsequent oncogenic mutations to arise on ecDNA such as *EGFRvIII*.

Third, it reveals the ability of ecDNA to co-amplify multiple weaker oncogenes to drive tumorigenesis, especially if they are located relatively close to each other in the chromosomal site of origin. Tumors with ecDNA amplified *PDGFRA* co-amplify other oncogenes on the same ecDNA and show different evolutionary trajectories compared to tumors with ecDNA amplified *EGFR* and no other co-amplified oncogenes.

Lastly, the development of our SPECIES model, based on both DNA, RNA, and spatial features, reveals that spatial-temporal evolutionary modelling can lead to unique insight into ecDNA- mediated oncogenesis. Intratumor heterogeneity of *EGFRvIII* contributes to failure of EGFR targeted therapies [37, 38], although CAR-T cells targeted both *EGFRvIII* and *EGFRwt* have recently shown promise[39]. The observation that amplification of wild-type *EGFR* on ecDNA, followed by mutational processes that give rise to more potent *EGFR* variants such as *EGFRvIII*, suggests that there may be a period between initial ecDNA formation and development of more potent gain-of-function mutagenesis, that could be amenable to therapeutic intervention and early cancer detection, for example through liquid biopsy which is under experimental investigation in brain tumors ([40, 41]. Several of the oncogenes amplified on ecDNA have been the subject of targeted therapy trials either alone or in combination, including *MET*, *CDK4*, *EGFR*, and *PDGFRA,* and other oncogenes are under experimental investigation such as *CLOCK, SOX13* and *AKT1* [42, 43]. It will be important to determine whether tumors with and without pre-malignant ecDNA accumulation respond differently to targeted therapies in such trials, and ecDNA characterisation may allow such molecular stratification (Fig 4j).

## Methods

### Patient Cohort

All 59 consecutive IDH1-wildtype adult glioblastoma cases surgically treated at a single UK center (University Hospital Southampton, UK) between March 2017 and June 2020 were included in this cohort. Full ethical approval was granted through Brain UK for this study. We obtained surgical tissue specimens (formalin fixed paraffin embedded, FFPE) from the clinical archives of the Department of Cellular Pathology, University Hospital Southampton. Samples were obtained from routine surgical resection, prior to starting chemotherapy or radiotherapy. IDH status was determined in routine clinical diagnostics, with immunohistochemistry using the IDH1 R132H mutation-specific antibody, supplemented with IDH1 / IDH2 sequencing as appropriate. All tumors retained ATRX expression.

The mean age of these patients was 61.6 years (range 38 - 80 years), of which 36 were male and 23 were female; 32 tumors displayed *MGMT* methylation (known to increase sensitivity to chemotherapy, [44]).

The FFPE specimens were used to generate tissue microarray (TMA) blocks. For each tumor sample, three regions were selected for TMA construction by a consultant neuropathologist including 1) tumor core, representing a solid tumor and avoiding as far as possible tumor necrosis and microvascular proliferation; 2) infiltrating zone, where tumor cells are invading into brain parenchyma, which retains its essential structure; and 3) leading edge, representing the most ‘normal’ brain tissue in the specimen, mostly comprising cortical grey matter and lacking significant numbers of morphologically detectable tumor cells (*n* = 177 samples). Two independent neuropathologists confirmed differentiation between the infiltrating zone and leading edge based on histology [17].

### Whole genome sequencing

Sequencing was performed using FFPE glioblastoma samples where tissue was available; a single 20um tissue section was used for DNA extraction using the Qiagen QIAmp DNA FFPE Advanced kit. Genomic DNA was randomly sheared into shorter DNA fragments, which were then end-repaired, A-tailed and further ligated with Illumina adapters. These fragments with adapters were size-selected, PCR-amplified and purified. Libraries were wantified through Qubit and qPCR; quantified libraries were pooled and sequenced by Illumina Novaseq 6000 (150 bp - paired end reads) with 15-fold coverage per tumor.

### ecDNA Detection and Characterisation

After aligning BAM files to GRCh38, CNVkit was used to detect DNA copy number alterations. Candidate seed regions with a copy number greater than 3.5 and size more than 20 kbp were identified using the AmpliconSuite-pipeline [18], which were leveraged for ecDNA characterisation with AmpliconArchitect and AmpliconClassifier. CycleViz (https://github.com/AmpliconSuite/CycleViz) enabled the visualization of candidate circular ecDNA structures. Oncogenes were classified based on the ONGene database [45], as well as glioblastoma driver genes previously reported (TCGA[46]).

### Nascent RNAscope and *EGFRvIII* Immunohistochemistry

We developed a method to detect nascent transcripts with RNAscope (which we term ‘nascent RNAscope’), based on probes for intronic sequences on *EGFR* / *PDGFRA* / *CDK4*. Although intronic probes have been used for ecDNA transcript localization with RNA FISH previously [29], use of probes for RNAscope facilitate multiplexed imaging. Multiplex RNAscope was run on the Bond RX using reagents and the standardized protocol from ACD Biotechne; RNAscope probes for EGFR, CDK4 and PDGFRA were designed against intronic gene sequences. The slides were imaged on the Akoya Vectra polaris multispectral slide scanner. This revealed high variability in expression of these oncogenes from cell-to-cell, reflecting the variability in copy number of these oncogenes (Supplementary Figure 19).

The FFPE slides were dewaxed and rehydrated to water through 100% and 70% ethanol. They were blocked with 0.3% hydrogen peroxide and then 5% BSA. The primary antibody (EGFRvIII Ab (D6T2Q) XP® Rabbit mAb, Cell Signaling Technology) was incubated at room temperature for 1 hour, then an anti-rabbit polymer was added for 1 hour. Then a DAB substrate was added for 10 minutes. The slides were then counterstained with haematoxylin and dehydrated through alcohols and a cover slip was applied.

### DNA FISH and Microscopy

Fluorescent *In Situ* Hybridization (FISH) was performed on 4uM Formalin Fixed Paraffin Embedded (FFPE) tissue sections according to a combination of the Agilent Technologies Protocol (Histology FISH Accessory Kit-K5799) and the Abbott Molecular Diagnostic FISH probe protocol. Briefly, FFPE sections were dewaxed in Xylene for 5 minutes followed by rehydration in 100%, 80% and 70% ethanol and then washed twice with Agilent Technologies wash buffer. The FFPE tissue was then incubated at 98°C for 10 minutes in Agilent Technologies pretreatment solution. The coplin jar is removed from the 98°C water bath and allowed to slowly cool for an additional 15 minutes. The FFPE slides were washed twice with Agilent Wash buffer. Stock pepsin (Agilent stock pepsin) was applied to the slide for 10 minutes at 37°C. FFPE slides were washed twice with Agilent wash buffer and then dehydrated using 70%, 80% and 100% ethanol prior to probe hybridisation. Gene Specific probes containing chromosome specific CEPs (Centromere Enumeration probes) against PDGFRA (Empire Genomics), EGFR (Vysis / Abbott) and MYC (Vysis / Abbott) were applied directly to the tissue sections with the coverslip being sealed with rubber solution glue.

Denaturation of the probes on the tissue was performed at 75^0^C for 7 minutes. The slide was then incubated overnight in a moist box at 37°C for 16 hours.

Slides were washed for 10 minutes at 73°C with 0.4X SSC containing 0.3% Igepal (Sigma) followed by a 10 minute room temperature wash with 2X SSC containing 0.1% Igepal. Slides are allowed to air dry and then counter stained with Vectashield mounting medium containing DAPI (ThermoFisher).

Images were captured using the VS200 Olympus Slide Scanner at 40x magnification. Quantification of FISH foci per nucleus was performed using QuPath software in a supervised fashion using the Cell Detection and Subcellular Cell Detection features, with the ‘includeClusters’ function to account for ecDNA clusters.

### Computational modeling of ecDNA driven tumors

We developed the SPECIES (*SP*atial *EC*dna Intratumor Evolution Simulation) framework to model expanding ecDNA driven tumor populations in a 2-dimensional space using an on-lattice cellular automaton model. Employing a kinetic Monte Carlo approach based on the method proposed by Bortz, Kalos and Lebowitz [47], and based on similar previous spatial agent-based models of tumor growth [48, 49], we simulate stochastic birth and death of tumor cells, as well as local cell displacement and random inheritance of ecDNA during cell division. Previous theoretical research into ecDNA dynamics in tumors supports the hypothesis of random segregation of ecDNA elements [15] described by a binomial partitioning process. We adopt the same approach to modelling ecDNA inheritance in our model. During cell replication, mother cells are able to displace neighboring cells in order to create the necessary space to divide.

See Supplementary Methods for full details on SPECIES. Code for simulating and plotting spatial tumors can be found at https://github.com/MagnusHaughey/spatial_ecDNA_patterns.

### Bayesian inference of patient tumor dynamics

We employ approximate Bayesian computation (ABC) [25,26] with rejection sampling to fit our computational model to individual patient data. Specifically, we estimate the values of model parameters *k*, *s* and *q* which best recapitulate the fraction of ecDNA-free tumor cells and the single-cell ecDNA copy number distribution from the tumor core and infiltrating margin of each patient tumor, derived using DNA FISH. To obtain spatial samples from each simulated tumor, we identify the core sample as the circular region of cells centered on the coordinates of the first tumor cell. The sampled ecDNA copy number distributions can vary considerably from one location to another at the tumor infiltrating margin. To capture this variability inherent to the system, we thus take 10 infiltrating margin samples from each simulated tumor, where each margin sample is taken as a circular region centered around a cell a radial distance of 75% of the tumor radius, at a random angle. We pair the core sample with each of the 10 infiltrating margin samples and treat these as 10 independent core-margin sample pairs, comparing each to the patient data. To avoid contamination with non-tumor cells, we filter out any single cells with an estimated ecDNA copy number of fewer than 3, both in the simulated and patient-derived samples. The size of the patient samples, after filtering, ranged between 583 to 6742 cells (median = 1396 cells), and we match the size of each sampled simulation core and margin region to the corresponding regions in each patient. We employ the Wasserstein metric [50, 51] to quantifiably compare the ecDNA copy number distributions in the simulated core and infiltrating margin samples to their patient-derived counterparts. To accept any given simulated core and infiltrating margin sample pair, require that they are both, individually, sufficiently near to their patient-derived counterparts. For purposes of summarizing the resulting model fit, we then sum the Wasserstein distances for core and infiltrating margin samples, to obtain the total distance between the simulation and patient data, denoted *σ*.

See Supplementary Methods for further details on model inference.

## Author contributions

I.N., M.H., C.S., & B.W. conceptualized this study. I.N. curated patient data. I.N. & M.H. performed formal analysis. I.N. & M.H. developed methodology. M.H. developed software. W.H., P.S.M., C.S. & B.W. supervised this work. I.N., M.H. & B.W. wrote the initial manuscript draft. All authors reviewed the manuscript.

## Acknowledgements

Tissue samples were obtained from BRAIN UK, which is supported by Brain Tumor Research and has been established with the support of the British Neuropathological Society and the Medical Research Council. I.N. is funded by a National Institute of Health Research Clinical Lectureship, CRUK Therapeutic Catalyst Award, University College London Biomedical Research Centre, the Academy of Medical Sciences and The Francis Crick Institute, which receives its core funding from Cancer Research UK (FC001169), the UK Medical Research Council (FC001169) and the Wellcome Trust (FC001169). C.S. is funded by CRUK (TRACERx (C11496/ A17786)), PEACE (C416/A21999) and CRUK Cancer Immunotherapy Catalyst Network), CRUK Lung Cancer Centre of Excellence (C11496/A30025), the Rosetrees Trust, Butterfield and Stoneygate Trusts, the NovoNordisk Foundation (ID16584), a Royal Society Professorship Enhancement Award(RP/EA/180007), the National Institute for Health Research (NIHR) University College London Hospitals Biomedical Research Centre, the CRUK–University College London Centre, the Experimental Cancer Medicine Centre, the Breast Cancer Research Foundation (US), and The Mark Foundation for Cancer Research Aspire Award (grant 21-029-ASP). J.L. and V.B. were supported by the National Cancer Institute (NCI) Informatics Technology for Cancer Research program (U24-CA264379). M.G.J. is supported by an NCI Pathway to Independence Award (NIH K99CA286968). This work was also supported by the Cancer Grand Challenges partnership funded by Cancer Research UK (CRUK) (P.S.M., CGCATF-2021/100012; V.B., and J.L., CGCATF-2021/100025) and the NCI (P.S.M., OT2CA278688; V.B., and J.L., OT2CA278635). P.S.M. is the eDyNAmiC team lead and V.B. is a member of the eDyNAmiC team. Work supervised by V.B. was also funded by The National Institutes of Health (NIH) grant R01- GM114362. B.W. is also supported by a Barts Charity Lectureship (grant no. MGU045) and a UKRI Future Leaders Fellowship (grant no. MR/V02342X/1).

