## Supplementary methods for "Extrachromosomal DNA driven oncogene spatial heterogeneity and evolution in glioblastoma"

**2 General model results 5**

**3 Parameter inference in human glioblastoma 7**

**4 Accuracy of ABC inference algorithm 9**

**5 Varying spatial location of core samples 10**

**6 Restricting parameter space of *k* 10**

**7 Varying cell birth/death ratio 11**

**8 Modelling ecDNA variant dynamics 11**

**9 Constant population size model of pre-expansion ecDNA dynamics 12**

**10 Modelling ecDNA co-amplification 13**

### 1 Spatial computational model of ecDNA-driven tumours

#### 1.1 Model initialisation

Each individual agent (*i.e.* tumour cell) in our model is defined by its (x,y)-coordinate in the lattice, and the number of ecDNA copies it is carrying. Each lattice position may either be unoccupied or occupied by strictly one tumour cell. We initiate an individual simulation by seeding a cell in the centre of a square lattice, carrying n copies of ecDNA. We specify the final size of the tumour in terms of number of cells, $N_{max}$, and estimate the approximate dimensions of the final system, assuming the final tumour to be perfectly circular, as $r = \sqrt{N_{max}/\pi}$ in units of lattice points. Independently of the value of ecDNA conferred replicative advantage, we assume the initial cell has already acquired the genetic alterations necessary for malignant growth and therefore has a birth/death ratio greater than 1.

After initialisation, we begin simulating stochastic cell birth and death until the tumour has reached a total size of $N_{max}$ cells. In its current form, the spatial model can only simulate tumours up to approximately $N_{max} = 10^6$ cells in a reasonable time-frame. Geometrically, if this were a cross-section of a 3D spherical tumour, lying on a plane containing the tumour centre, this would correspond to a 3D tumour of approximately $10^9$ cells (tumour volume of approximately 1 $cm^3$). If the patient tumours were significantly larger than $10^9$ cells at the point of resection, we may under-estimate the selection strength of ecDNA-carrying cells, or the number of ecDNA copies in the tumour initiating cell, by comparing them to 2D simulated tumours of size approximately equal to $10^6$ cells. For the current ABC inference, we simulated tumours of $10^6$ cells, and acknowledge the limitations this might impose on the inferences made in our patient tumours.

#### 1.2 Random inheritance of ecDNA

Inheritance patterns of ecDNAs were recently characterised in various tumour cell lines, and were well described by a binomial partitioning process [1]. In line with this evidence, we partition the ecDNA in a dividing mother cell into the two daughter cells in the following way: we assume each ecDNA element is replicated strictly once, and thus a mother cell containing $X$ copies of ecDNA results in daughter cell copy numbers, $x_1$ and $x_2$, of

$$x_1 \sim Binomial(2X, \frac{1}{2}), \tag{1}$$

$$x_2 = 2X - x_1. \tag{2}$$

#### 1.3 Cell displacement

Several approaches have been used in previous studies for implementing cell displacement in agent-based models, including non-linear cell pushing [2] mechanics and local cell motility [2–8]. Here, we model cell displacement using a linear cell pushing algorithm [8–12], which involves the creation of empty space in the neighbourhood of dividing cells by pushing other cells in a straight line, filling a distal empty lattice point.

We implement cell pushing using the following method. After a cell has been selected for division, we identify its nearest empty point on the lattice. If there are multiple nearest lattice points, then one is chosen at random with uniform probability. Next, by drawing a straight line between the dividing cell and this empty lattice point, we construct a path of cells connecting these two locations. Last, each cell within this path is moved one position along the path, towards the empty lattice point (the dividing cell is not moved during this process). This process results in the filling of the empty lattice point, and the creation of a new empty space in the immediate neighbourhood of the dividing cell, enabling it to divide into two daughter cells (one daughter cell occupies the old position of the parent cell, and the other is placed in the neighbouring empty lattice point).

Spatial structure of the tumour influences the pattern of evolution [5, 8], and so we explore different tumour growth regimes in our simulations, achieved by varying a model parameter $q$ which sets the range within which a dividing cell may search for a nearby empty lattice point. The value of $q$ is always a global parameter i.e. all cells in the system are subject to the same $q$ value. If there are no empty lattice points within a radius $q$ of the dividing cell, the cell will not divide but will remain in its current position. Model parameter $q$ can take any integer value greater than zero. Representing the strongest possible spatial constraints: when $q = 1$, dividing cells may only search for empty lattice points within a radius of one lattice unit (von Neumann neighbourhood). By setting $q$ to be very large, we are able to remove all spatial constraints (in this study, we typically set $q = 1000$ to represent this regime).

#### 1.4 Differential selection

We assume that presence of ecDNA can confer a positive selective advantage to a tumour cell, and implement this by means of an increase in replication rate. In general, it is possible to implement any mathematical function to describe ecDNA-conferred cell fitness, however we adopt a simple ecDNA copy number independent model, leading to a cell replication rate, $r_b$, of

$$r_b(x, s) = \begin{cases} 1 & x = 0 \\ 1 + s & x > 0 \end{cases} \tag{3}$$

where $x$ denotes the ecDNA copy number in the cell and s is a model parameter which specifies the strength of positive selection. Neutral selection, where ecDNA presence confers no replicative advantage to the cell, is achieved by setting $s = 0$, whereas $s > 0$ describes positive selection. Whilst it may be reasonable to expect the replication rate of a cell to depend more strongly on the number of ecDNA-amplified oncogenes, copy number independent selection models have been employed in other mathematical models [1, 13], and have been shown to adequately describe the dynamics of ecDNA driven tumours.

##### 1.5 Cell death

We model cell death as an independent event in the simulations, uncoupled from cell division. The rate of cell death, $r_d$, is fixed at some fraction of the neutral cell division rate, $r_b(x, 0)$,

$$r_d = r_b(x, 0) \cdot \alpha, \tag{4}$$

where $s$ denotes the strength of ecDNA conferred replicative advantage, $x$ represents the ecDNA copy number in the cell and $\alpha \in [0, 1]$, which was set at $\alpha = 0.5$ throughout this study unless stated otherwise. Following a cell death event, the dead cell is immediately removed from the lattice, leaving behind an empty lattice point.

##### 1.6 Stochastic algorithm

We adopt a kinetic Monte Carlo approach based on the method proposed by Bortz, Kalos and Lebowitz [14]. For each iteration of the algorithm, we first select the event to occur by considering the total sum of all event rates in the system, before choosing the cell in which the event will take place.

In our model we consider stochastic cell birth and death which, in a cell with $x$ copies of ecDNA, occur with rates $r_b(x, s)$ and $r_d$ respectively. As the rate of cell death does not depend on the ecDNA copy number, the total rate of cell death in the system, $R_d$ , is

$$R_d = N \cdot r_d, \tag{5}$$

where $N$ denotes the total number of cells present in the system. In its most general form, we partition cell birth into separate events for each value of ecDNA copy number, thus allowing for ecDNA copy number dependent selection functions. Denoting $N_x$ as the number of cells with $x$ copies of ecDNA, the sum of birth rates across all cells with copy number $x$, $R_b^x$, is given as

$$R_b^x = N_x \cdot r_b(x, s). \tag{6}$$

For each iteration of the algorithm, we therefore compute the sum of birth and death rates across all cells in the system, $R_{tot}$, as

$$R_{tot} = R_d + \sum_{x=0}^{x_{max}} R_b^x, \tag{7}$$

where $x_{max}$ denotes the largest single-cell ecDNA copy number in the system. To determine which event type will occur, we generate a uniform random variable $\alpha \in \text{Uniform}(0, 1)$ and select cell death if

$$\alpha \cdot R_{tot} < R_d. \tag{8}$$

Otherwise, division occurs for a single cell with $x$ ecDNA copies, for $x$ satisfying

$$\sum_{i=0}^{x-1} R_b^i < (\alpha \cdot R_{tot} - R_d) \leq \sum_{i=0}^{x} R_b^i. \tag{9}$$

Once the type of event has been determined, a cell to undergo the event must be chosen. We thus choose, with uniform probability, a single cell from the pool of all eligible cells (*e.g.* if the event is division of a cell with $x$ copies of ecDNA, then all cells in the system containing $x$ ecDNA are eligible to be chosen).

For the majority of the modelling performed in this study, we employed a simple ecDNA copy number independent model of selection. This greatly simplifies the process of selecting events since all ecDNA copy number states of $x > 0$ can be combined into one single group with combined rate, denoted $R_b^{x>0}$, given as

$$R_b^{x>0} = N_{x>0} \cdot r_b(x > 0, s), \tag{10}$$

where $N_{x>0}$ denotes the number of cells containing one or more copies of ecDNA.

### 2 General model results

We implemented the spatial computational model set out above to study the effects of initial ecDNA copy number, $k$, ecDNA conferred selection strength, $s$, and the strength of spatial constraints, $q$, on the resulting spatial patterns of ecDNA copy number in the expanded tumour (Supplementary Figure 5). When model parameter $q$ is sufficiently large ($q = 1000$ in our implementation) all spatial constraints are removed and our model recapitulates previously published results, derived using non-spatial models, describing the time evolution of the single-cell ecDNA copy number distribution and the fraction of tumour cells carrying zero copies of ecDNA [1] (Supplementary Figure 20a).

Our model builds upon previously studied models not only by explicitly including spatial constraints, but also by allowing for different numbers of ecDNA copies in the initial tumour cell. For tumours with no spatial constraints, both small and large values of initial cell ecDNA copy number, $k$, lead to a mean single-cell ecDNA copy number which is spatially homogeneous across the entire tumour (Supplementary

Figure 21). When spatial constraints are included, mean copy number is either constant with increasing radial distance from the tumour centre when ecDNA confers no selective advantage ($s = 0$) or increasing approximately linearly when ecDNA-positive cells are under positive selection ($s > 0$). The mode of growth of the tumour and the initial ecDNA copy number also affect the variation in ecDNA copy number at the single cell level. For non-spatially constrained tumours, the standard deviation of single cell ecDNA copy number is roughly constant, or weakly increasing, with radial distance from the tumour centre (Supplementary Figure 22). Conversely, for tumours with strong spatial constraints, variation in single-cell copy number increases more markedly as cells are sampled further away from the tumour centre. For $k = 1$, the standard deviation is lower for neutral dynamics, compared to cases of positive selection, both for weak and strong spatial constraints. Interestingly, this is reversed for larger initial ecDNA copy numbers, with neutral dynamics leading to larger standard deviation. This is likely a result of the ecDNA-negative state being an absorbing state: once a cell has lost all its copies of ecDNA, it may never regain them (since we do not model *de novo* ecDNA production). For tumours which begin a cell with a single copy of ecDNA, many resulting descendant cell lineages will be stuck in the ecDNA-negative state, driving down the overall variation in ecDNA copy number at the single-cell level. When tumours begin with many copies of ecDNA, neutral dynamics lead to larger variation as fewer cells are ecDNA-negative. When $s > 0$, however, those ecDNA-negative cells which arise in the tumour are selected against, and thus their relative fraction within the tumour population is reduced, driving down the variation at the single-cell level.

The initial number of ecDNA copies has a large impact on the probability of the tumour retaining or losing ecDNA during expansion. Simulations of tumour growth from a single cell revealed that a starting ecDNA copy number of 1 led to complete loss of ecDNA in more than 70% of fully formed tumours in the absence of ecDNA conferred fitness advantages, with many tumours also failing to maintain ecDNA populations even when ecDNA give rise to a positive selective advantage. In contrast, presence of 20 ecDNA copies in the initiating cell resulted in near ubiquitous maintenance of ecDNA during tumour formation (Supplementary Figure 23).

The impact of including spatial constraints are most pertinent when one compares the overall single-cell ecDNA copy number distribution for tumours with and without spatial constraints (Supplementary Figure 20b). Our model shows that including spatial constraints leads to a wider overall distribution, with more cells at both low and high extremes of ecDNA copy number, with the exception of when $s = 0$ and $k = 1$. This increased variation, under-represented by previous non-spatial models, could increase the ability of the tumour to adapt quickly to changing selective pressures, such as introduction of chemotherapy, targeted therapies or radiotherapy.

The underlying source of the increased variation for tumours with and without spatial constraints can be understood by comparing the distribution of cell lineage lengths in each case (Supplementary Figures

20c-d). When there are no spatial constraints, all cells are equally capable of dividing (under neutral dynamics) and thus the distribution of cell lineages in the final tumour is tightly peaked around a mean value (theoretical mean for a tumour of size $N_{max} = 10^5$ cells is $\log_2(10^5) \approx 16.6$, however for our simulations this mean was marginally greater as a result of our assumption that cell cycle times are exponentially distributed). Spatial constraints during tumour growth lead to surface driven dynamics, whereby only cells situated at the expanding edge of the tumour are capable of division. This leads to a spatial bottleneck, resulting in some cell lineages (those surfing at the leading edge) to be significantly greater than others (*e.g.* those for cells trapped within the interior of the tumour) and a larger overall mean cell lineage length. Since ecDNA is binomially distributed at each cell division, increasing the variation in single-cell ecDNA copy number in each instance, tumours with stronger spatial constraints, and therefore a higher mean cell lineage length, will have a wider and more varied distribution of ecDNA copy numbers.

### 3 Parameter inference in human glioblastoma

We employ approximate Bayesian computation (ABC) [15,16] with rejection sampling to fit our computational model to individual patient data. Our method of parameter inference can be summarised in the following steps:

1. Define prior distributions for model parameters $k$, $s$ and $q$.
2. Sample candidate parameter values from prior distributions, $k^*$, $s^*$ and $q^*$, and execute a spatial simulation with these model values as input parameters.
3. Sample regions of simulated tumour from the tumour core and infiltrating margin, and derive single-cell copy number distributions for both sampled regions.
4. Quantitatively compare both patient single-cell copy number distributions and fraction of ecDNA-free tumour cells from tumour core and infiltrating margin to their simulated counterparts.
5. If simulated data is sufficiently similar to patient data, add parameter values $k^*$, $s^*$ and $q^*$ to posterior sample set.
6. Return to step 2 and repeat until the posterior sample set is of size $N$.

For the parameter estimation performed in this study, we specify prior distributions $P(k)$, $P(s)$ and $P(q)$ of

$$P(k) = \text{Uniform}(1, 150), \tag{11}$$

$$P(s) = \text{Uniform}(0, 5), \tag{12}$$

$$P(q) = 1, 2, 5, 10, 50, 1000. \tag{13}$$

To obtain spatial samples from simulated tumours, we identify the core region as the circular region of cells centred on the coordinates of the first tumour cell. We sample 10 infiltrating margin samples from each simulated tumour, each taken as circular regions centred around a cell a distance of 75% of the tumour radius, each at a random angle. We pair the core sample with each of the 10 infiltrating margin samples and treat these as independent samples when comparing simulated to patient data. Additionally, we match the size of each sampled core and infiltrating margin region to the corresponding regions in each patient.

Prior to comparing simulated and patient data, we filter all patient core and infiltrating margin samples to remove any cells with an estimated ecDNA copy number fewer than three (referred to as low-ecDNA cell fraction herein). This is because our estimates of ecDNA copy number in this range, derived using image analysis software applied to DNA FISH images (cross reference to methods section about DNA FISH), are particularly noisy. Furthermore, from DNA FISH images alone, we are unable to distinguish between non-tumour cells and tumour cells which are carrying zero copies of ecDNA. Knowledge of the low-ecDNA cell fraction is, however, an important indicator of the underlying tumour dynamics. To exploit this information, we record the fraction of tumour cells removed in the patient data during filtering, and use these measurements as additional data with which to fit patient to simulated data.

To quantify the similarity of simulated and patient derived ecDNA copy number distributions, we thus combine the use of two metrics. First, we employ the Wasserstein distance [17, 18], a commonly used metric in statistical inference and machine learning. During the data comparison stage of our ABC algorithm (step 5 above), we compute the Wasserstein distance between simulated and patient ecDNA copy number distributions, both for the tumour core and infiltrating margin, and denote these distances as $\delta_1^c$ and $\delta_1^m$ respectively. Second, we compare the low-ecDNA cell fraction for the patient and simulated data, both for the core and infiltrating margin regions, using the expression

$$\delta_2 = \frac{|n_{sim} - n_{pat}|}{N_{pat}}, \tag{14}$$

where $n_{sim}$ and $n_{pat}$ denote, respectively, the sample size in the simulated and patient data after filtering low-ecDNA cells, and $N_{pat}$ denotes the sample size in the patient prior to filtering. We compute this comparison metric both for the core and margin regions, and denote these as $\delta_2^c$ and $\delta_2^m$ respectively. In order to accept a simulation into the posterior sample set, we require that each simulation/patient comparison metric must be smaller than some threshold value, *i.e.*

$$\delta_1^c < \epsilon_1^c, \tag{15}$$

$$\delta_1^m < \epsilon_1^m, \tag{16}$$

$$\delta_2^c < \epsilon_2^c, \tag{17}$$

$$\delta_2^m < \epsilon_2^m, \tag{18}$$

where threshold values on low-ecDNA cell fraction similarity, $\epsilon_2^c$ and $\epsilon_2^m$, were chosen so as to optimise the accuracy of the inference algorithm. Threshold values on the Wasserstein distances, $\epsilon_1^c$ and $\epsilon_1^m$, were determined on a patient-specific basis such that the ABC inference algorithm achieved an acceptance rate of $\leq 5\%$.

After a sufficiently large number of repetitions, we obtain a good approximation for the posterior distribution of ($k$, $s$, $q$) which represents the regions of the parameter space for which there is most model support. From these distributions, we estimate the optimal parameter values by measuring the median of the 1-dimensional marginal distributions for each of $k$, $s$ and $q$. We estimate errors as the variance of these marginal distributions.

### 4 Accuracy of ABC inference algorithm

We explored the sensitivity and specificity of our model inference approach by applying it to a large set of simulated tumour core and infiltrating margin pairs, generated using the computational model itself. We generated an artificial patient dataset consisting of 100 simulated tumours for each combination of $k \in \{1, 25, 50, 100\}$, $s \in \{0, 0.5, 1, 2\}$ and $q \in \{1, 5, 50, 1000\}$ (Supplementary Figure 24). From each simulated tumour, we extracted a sample of tissue from the tumour core and from the outer edge at a randomly chosen angle. All tumours were simulated up to a final size of $10^5$ cells and sampled regions were fixed at a size of 5000 cells.

To determine the optimal values for thresholds ($\epsilon_1^c$, $\epsilon_1^m$, $\epsilon_2^c$, $\epsilon_2^m$), we computed the sensitivity and specificity (true positive and true negative rate respectively) across the range of parameter values represented in our artificial patient dataset, repeating this for a range of threshold values. We selected the combination of threshold values which maximised the sensitivity and specificity of the algorithm (Supplementary Figure 25a).

Results from this test demonstrate that the inferences made by the algorithm are reasonably accurate across all parameter values represented in the artificial patient dataset (Supplementary Figure 25b). Sensitivity and specificity of the algorithm is particularly high across the full range of $k$ and $q$ values we tested, with the algorithm particularly able to recover the true underlying $k$ value in the patient data. Due to our use of a constant selection model in our simulations, we were less successful at determining the true underlying selection coefficient, $s$. This loss of sensitivity is likely due to the fact that in tumours with a high fraction of ecDNA-positive cells, neutral competition will mostly ensue regardless of the specific value of $s$. Despite this, the inference algorithm still displayed moderate sensitivity when we pool all positive selection coefficients and simply predict neutral ($s = 0$) or positive ($s > 0$) selection. A prediction of neutral selection is made if the error on the point estimator for the selection coefficient, $s$, encompasses $s = 0$, otherwise the sample is classed as being consistent with positive selection.

### 5 Varying spatial location of core samples

When sub-sampling tissue from our simulated tumours, the core sample is always centred around lattice coordinates of the tumour initiating cell, however the same regularity cannot be obtained when sampling tissue from real tumours. To understand how this aspect of the model affects our measured ecDNA copy number distributions, and thus our inferred model parameters, we employed an alternative tissue sampling scheme, in which we sampled tissue from a region a short radial distance from the coordinates of the tumour initiating cell, at a random angle. We repeated the parameter inference using this alternative approach, but found the same oncogene-level patterns of inferred parameters, suggesting that variations in core sample location is not responsible for this observed pattern.

### 6 Restricting parameter space of $k$

To further investigate the model prediction in some patients of high initial ecDNA copy number, $k$, we repeated the parameter inference whilst constricting the value of $k$ to $k = 1$ only (Supplementary Figure 15). For each patient, we determined the optimal model fit to the ecDNA copy number distribution and computed the difference in affinity between model and data, denoted $\Delta\sigma$, for the normal parameter inference and the n-restricted case. We found that the model best-fit in the $k$-restricted inference was notably worse (indicated by a large $\Delta\sigma$) in patients predicted to have a high $k$ in our initial parameter inference. Whilst this does not rule out alternative explanations such as copy number dependent selection or co-selection/co-segregation dynamics, this supports the notion that, within the context of our model, a high value of $k$ is the only sufficient explanation for the observed ecDNA copy number distributions in some of our patients.

### 7 Varying cell birth/death ratio

Throughout all model simulations in this study, we maintained a constant cell death/birth ratio of $\psi = 0.5$. To test the sensitivity of our patient inferences on this parameter, we repeated the patient model fitting with death/birth ratios of $\psi = 0.1$, leading to few cell deaths during expansion, and $\psi = 0.9$, giving rise to near-balanced cell birth and death (Supplementary Figure 14). Inferred model parameters for each patient were very similar under these two alternative regimes to the original inferred values using $\psi = 0.5$, suggesting weak sensitivity to this model parameter. Inferences for model parameter $k$ were particularly robust to these changes. Reducing the death/birth ratio to $\psi = 0.1$ slightly reduced the inferred ecDNA selection strength, $s$, and cell pushing strength, $q$, whilst an increased value of $\psi$ led to slightly greater inferred values of $s$ and $q$.

### 8 Modelling ecDNA variant dynamics

Both structural variant and immunohistochemistry analysis confirmed the presence of *EGFR* mutations amplified on ecDNA *in vivo*. We modelled the initial dynamics of wild-type ecDNA, followed by the emergence of an advantageous variant of the gene on ecDNA within the framework of SPECIES. To explore the dynamics of the wild-type and variant ecDNA, we varied the following model parameters: $k_{wt} > 0$ and $k_{var} \geq 0$, which respectively set the number of wild-type and variant ecDNA copies in the clone-initiating cell; and $V_{var} \geq 0$, which specifies the the size to which the tumour must expand (in terms of total number cells) before the mutant ecDNA emerges from within the existing wild-type ecDNA population. Setting $V_{var} = 0$ gives rise to a tumour with "pre-expansion" mutant ecDNA, the number of which in the clone-initiating cell is specified by $k_{var}$, whereas $V_{var} > 0$ leads to a tumour with "post-expansion" mutant ecDNA.

We model evolutionary dynamics by assuming that wild-type and variant ecDNA confer a replicative advantage to tumour cells, with associated selection coefficients denoted $s_{wt}$ and $s_{var}$ respectively, with $s_{wt} \leq s_{var}$. Specifically, a tumour cell with $x_{wt}$ copies of the wild-type ecDNA and $x_{var}$ copies of variant ecDNA will have a replicative rate, $r_b$, given by

$$r_b(x_{wt}, x_{var}, s_{wt}, s_{var}) = \begin{cases} 1 & (x_{wt} = 0 \quad \text{and} \quad x_{var} = 0), \\ 1 + s_{wt} & (x_{wt} > 0 \quad \text{and} \quad x_{var} = 0), \\ 1 + s_{var} & (x_{var} > 0), \end{cases} \tag{19}$$

where we assumed that the wild-type ecDNA conferred a weakly positive advantage, $s_{wt} = 0.2$, and the mutated ecDNA conferred a stronger advantage, $s_{var} = 2$. We further assume that each ecDNA is replicated once during cell division, and that all ecDNAs are partitioned binomially across the two resulting daughter cells. We set the final simulated tumour size to be $N_{max} = 10^5$.

Using this model, we computed the mean measured ecDNA heteroplasmy (percentage of ecDNA copies in sample carrying wild-type version of the gene) in the tumour core for each combination of $k_{wt} \in \{1, 10, 50\}$ and $k_{var} \in \{1, 10, 50\}$ for pre-expansion ecDNA mutation ($V_{var} = 0$), and $k_{wt} \in \{1, 10, 50\}$ and $V_{var} \in \{10^2, 10^3, 10^4\}$ for post-expansion ecDNA mutation ($k_{var} = 0$) (Supplementary Figures 26 & 27). We simulated 1,000 tumours for each parameter combination, and computed the mean core heteroplasmy for each combination of parameters. To measure core heteroplasmy, we extracted 2,000 cells from the core of each simulated tumour and computed the ecDNA heteroplasmy as $X_{wt}/(X_{wt} + X_{var})$, where $X_{wt}$ and $X_{var}$ denote the total number of wild-type and variant ecDNA copies, pooled across the entire sampled set of cells. These circular core samples were taken a radial short distance from the coordinates of the tumour initiating cell, at a random angle, in order to emulate the variation in sample location in the patients' samples.

Alongside measuring ecDNA heteroplasmy in the tumour core, we computed the probability of observing any variant ecDNA in randomly sampled regions of the tumour core and infiltrating margin (Supplementary Figure 28). For the same combinations of parameters set out above, we simulated 1,000 tumours, sampling 5 core & margin pairs from each tumour, with each regional sample consisting of 2,000 cells.

### 9 Constant population size model of pre-expansion ecDNA dynamics

To further understand the dynamics of wild-type and variant ecDNA in the pre-malignant tissue, we exploited the simulation framework of SPECIES to model a non-spatial, constant population size cell population. These simulations follow the dynamics between the generation of the first wild-type ecDNA up to the first mutation of an ecDNA. After the system of cells is initialised (fixed at 1,000 cells), one cell in the system is selected randomly to acquire a single copy of the wild-type ecDNA. The system is then evolved following a Moran process [19], assuming the same dynamics of ecDNA as the original SPECIES simulations (ecDNA replication and random inheritance). The model of ecDNA selection is also the same as the original SPECIES model, with the wild-type ecDNA conferring a weakly positive advantage to the host cell ($s = 0.2$). During cell division, each copy of wild-type ecDNA in the mother cell is replicated once, and may stochastically mutate to the variant ecDNA with rate $\mu$ per ecDNA replication. We simulate the system up until the first ecDNA mutation, at which point we measure the ecDNA heteroplasmy of the cell in which the mutation occurred.

#### 10 Modelling ecDNA co-amplification

Whole-genome sequencing of the core tumour samples shed further light on the extent of ecDNA co-amplification in *in vivo* GBM. Of the 57 tumour cores we sequenced, we found 8 which contained multiple oncogene-amplifying ecDNA amplicons. Recently co-segregation (correlated inheritance) and co-selection (selective advantage for maintaining a mixture of both ecDNA species within a cell) has been shown to play a role in maintaining ecDNA populations in tumour cell lines [13]. Our computational model only accounts for a single ecDNA type, however, and thus does not capture these important dynamics in our patient-derived samples. In particular, it may be that co-segregation and co-selection dynamics could provide an alternative explanation for a subset of patient samples which were previously consistent with a large number of pre-existing ecDNA copies (*i.e.* high inferred $k$).

To explore this, we adapted SPECIES to include two ecDNA types. We implemented co-segregation dynamics using the same approach as Hung et al. [13] and extended our constant selection model to account for both types of ecDNA. In this adapted model, a tumour cell with $x_1$ copies of ecDNA type 1 and $x_2$ copies of ecDNA type 2 will have a replicative rate, $r_b$, given by

$$r_b(x_1, x_2, s_p, s_m) = \begin{cases} 1 & (x_1 = 0 \quad \text{and} \quad x_2 = 0), \\ 1 + s_p & (x_1 > 0 \quad \text{and} \quad x_2 = 0), \\ 1 + s_p & (x_1 = 0 \quad \text{and} \quad x_2 > 0), \\ 1 + s_p + s_m & (x_1 > 0 \quad \text{and} \quad x_2 > 0), \end{cases} \tag{20}$$

where $s_p \geq 0$ denotes the strength of positive selection for the "pure" state *i.e.* when only 1 type of ecDNA is present, and $s_m \geq 0$ for the "mixed" state *i.e.* containing both types of ecDNA.

In principle one could obtain ecDNA copy number distributions for both ecDNA types using this alternative model, however we had only single-cell resolution copy number data for one of the two ecDNA amplicons in each patient (with the exception of patient A5, for whom we had data both for the EGFR- and PDGFRA-amplifying ecDNA amplicons). We thus sampled data for one ecDNA type when fitting the co-amplified ecDNA model to the patient data, however this meant we were unable to reliably infer specific values co-segregation and co-selection.

#### 11 Estimation of DNA-RNA mapping on ecDNA

To understand the relationship between ecDNA copies and transcription, we developed a computational tool that finds an optimised scaling parameter to infer the correlation between the ecDNA copies and the expression levels. We implemented a Monte Carlo inference which adds a decreasing exponential noise,parameterised by $\lambda$,such that the distance between the nascent RNAScope and the DNA-FISH count distributions for a given patient sample is minimised. The framework starts by removing cells with 0, 1 and 2 ecDNA copies, to take into account the uncertainty of quantifying ecDNAs from FISH in cells with few copies. We then sample a random value $\lambda_i$ from a uniform distribution Uniform$(0.1, 4)$ and reconstruct a scaled DNA-FISH counts $k_i'$ by sampling from an exponential decreasing probability function for every ecDNA copy $k$ found in the original DNA-FISH distribution, *i.e.*

$$k_i' \sim \text{Exp}(-\lambda i k). \tag{21}$$

We compute the distance between the scaled DNA distribution, $x_i$, and the RNAScope distribution, $y$, using three distances, $E(x_i, y)$, $M(x_i, y)$ and $K(x_i, y)$, where $E$ and $M$ are relative Euclidean distance for the entropy and the mean of the distributions respectively, and $K$ is the Kolmogorov-Smirnov statistic. These distances are grouped into a vector of summary statistics $S(x_i, y) = [E(x_i, y), M(x_i, y), K(x_i, y)]$. We tested a total of $100,000$ random $\lambda_i$ values for each patient, and constructed the posterior distribution of $\lambda$ using the 500 values which give rise to the smallest $S(x_i, y)$. We then measure the modal value of the posterior distribution to find the optimal scale parameter $\lambda^*$. In Fig. 19, we plotted for all patients the Wasserstein distance between the matched DNA-FISH counts scaled by $\lambda^*$ and the RNAScope distributions.
