## Supplementary figures for "Extrachromosomal DNA driven oncogene spatial heterogeneity and evolution in glioblastoma"

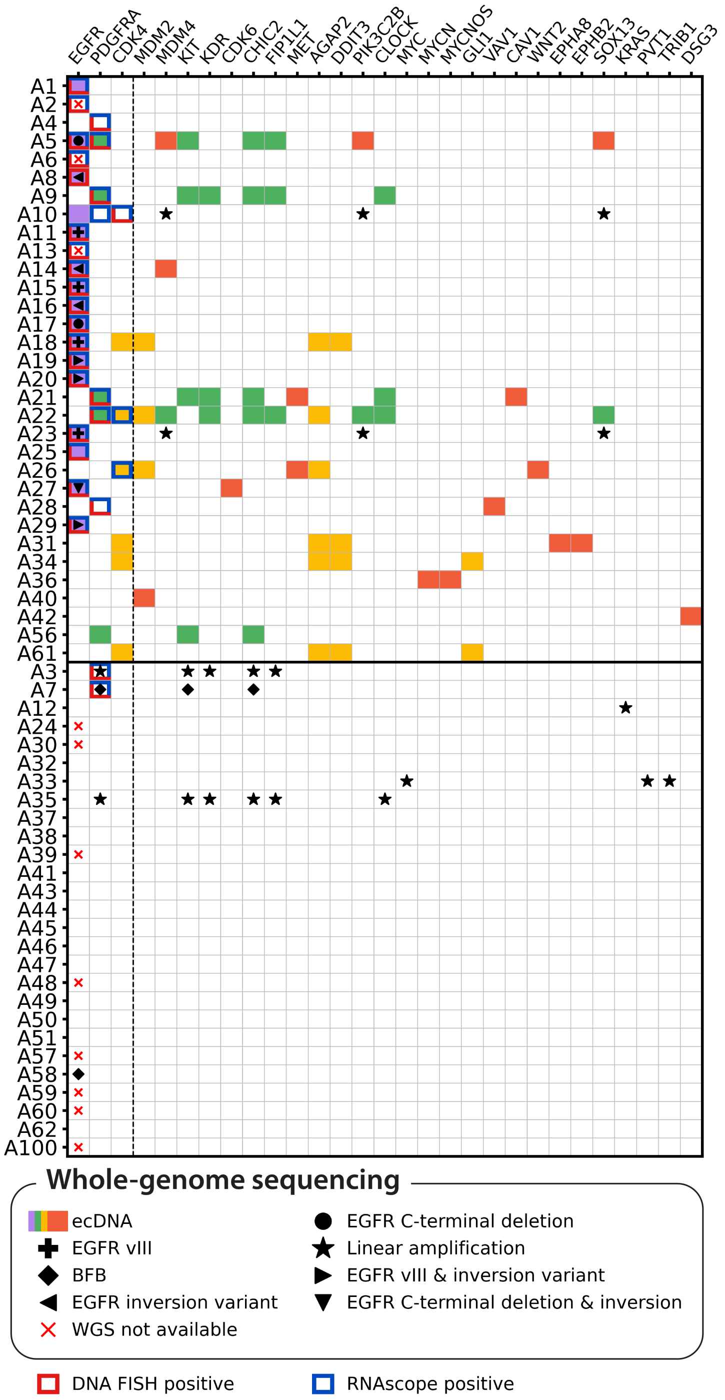


**Supplementary Figure 1:** Waterfall plot showing focal copy number amplifications across the GB-UK cohort ($n = 59$ patients), categorised by amplicon type (ecDNA, breakage-fusion-bridge (BFB) cycle, linear). Horizontal line between patients A61 & A3 separate patient samples with (above) or without (below) ecDNA, detected either by whole-genome sequencing, DNA FISH or nascent RNAscope. Oncogenes tested with DNA FISH and / or nascent RNAscope, *EGFR*, *PDGFRA* and *CDK4* are separated to the left of the vertical dashed line.

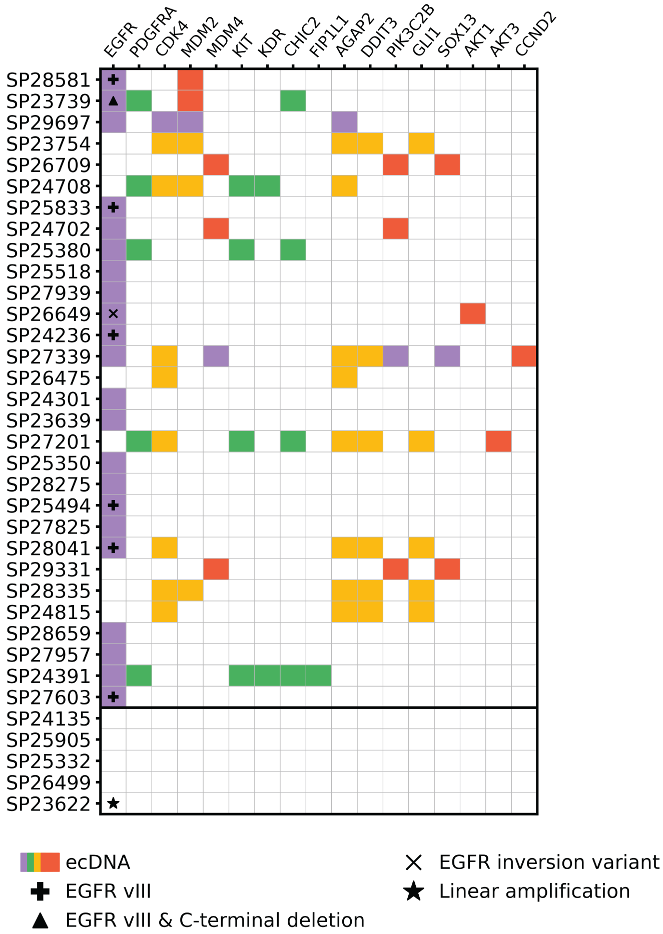


**Supplementary Figure 2:** Waterfall plot showing focal copy number amplifications across the PCAWG cohort ($n = 35$ patients), categorised by amplicon type (ecDNA, breakage-fusion-bridge (BFB) cycle, linear) Horizontal line between patients SP27603 & SP24135 separate patient samples with (above) or without (below) ecDNA, detected either by whole-genome sequencing, DNA FISH or nascent RNAscope. Oncogenes tested with DNA FISH and / or nascent RNAscope, *EGFR*, *PDGFRA* and *CDK4* are separated to the left of the vertical dashed line.

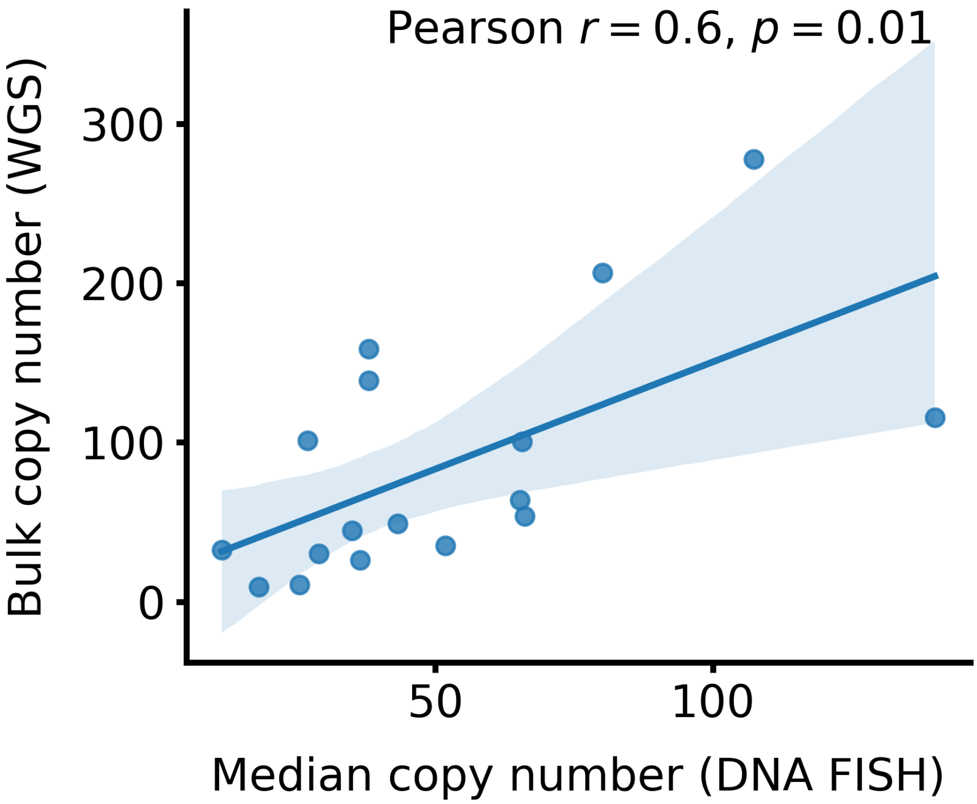


**Supplementary Figure 3:** Correlation between measured ecDNA oncogene copy numbers, measured using DNA FISH and whole-genome sequencing.

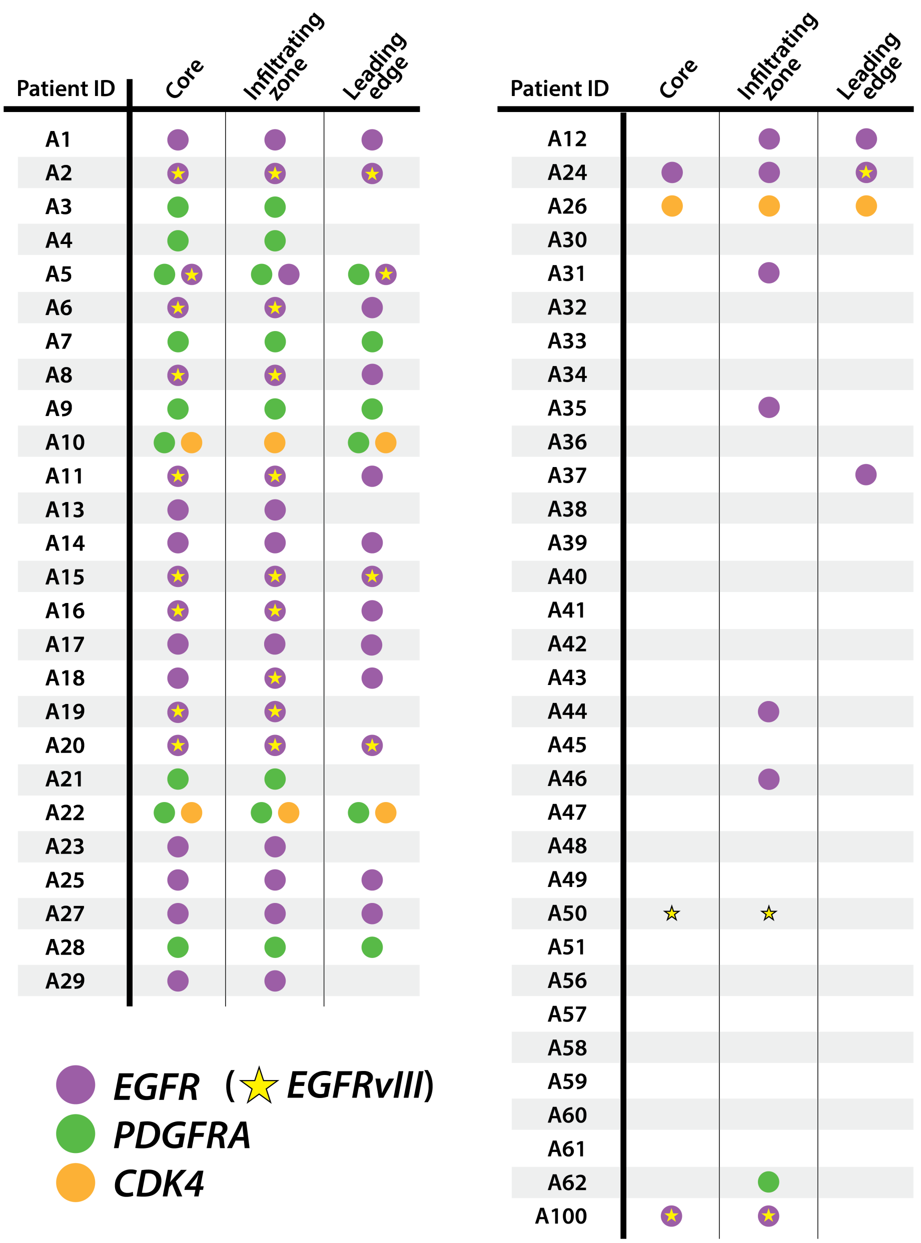


**Supplementary Figure 4:** Oncogenic ecDNA observations in tumour core, infiltrating zone and leading edge locations across entire patient cohort. Data represent combined DNA FISH and RNAscope observations

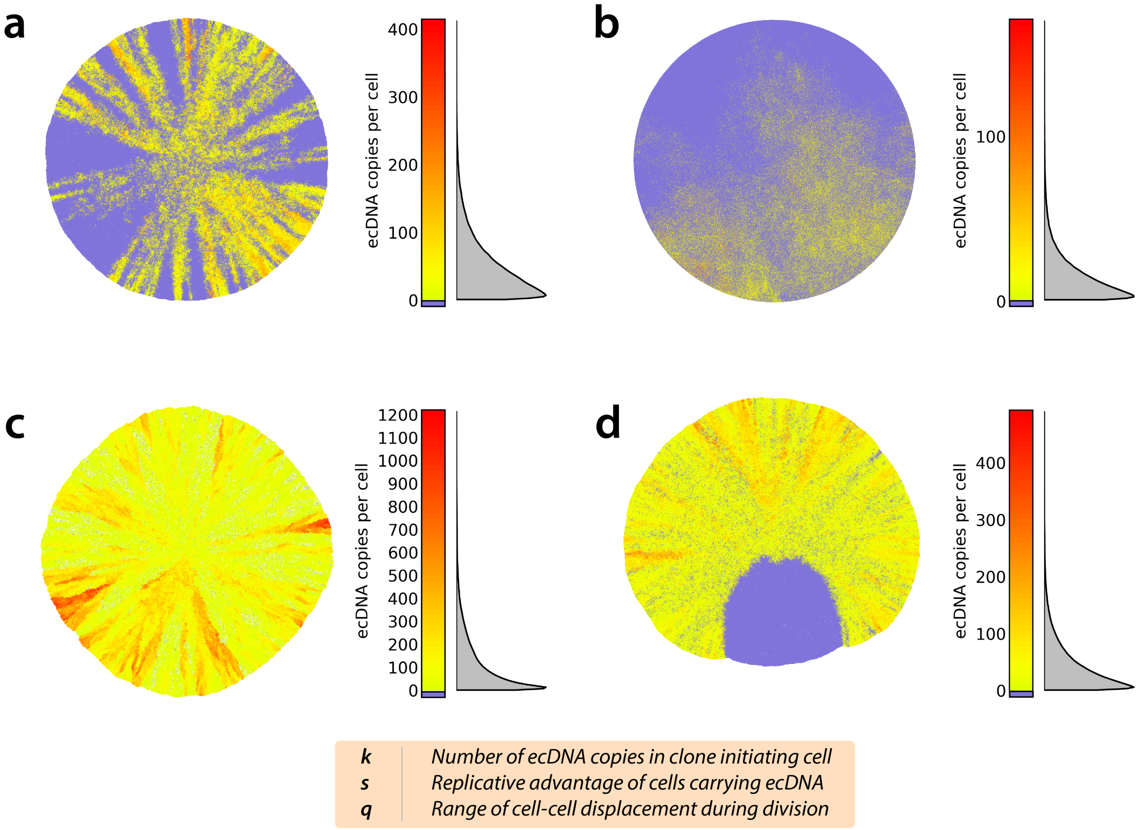


**Supplementary Figure 5:** Example ecDNA-driven tumours simulated with SPECIES for **(a)** $k = 16$, $s = 0$, $q = 10$; **(b)** $k = 15$, $s = 0$, $q = 1000$; **(c)** $k = 105$, $s = 4.9$, $q = 2$ and **(d)** $k = 1$, $s = 0.2$, $q = 10$, showing typical range of simulated tumour ecDNA patterns attainable with SPECIES.

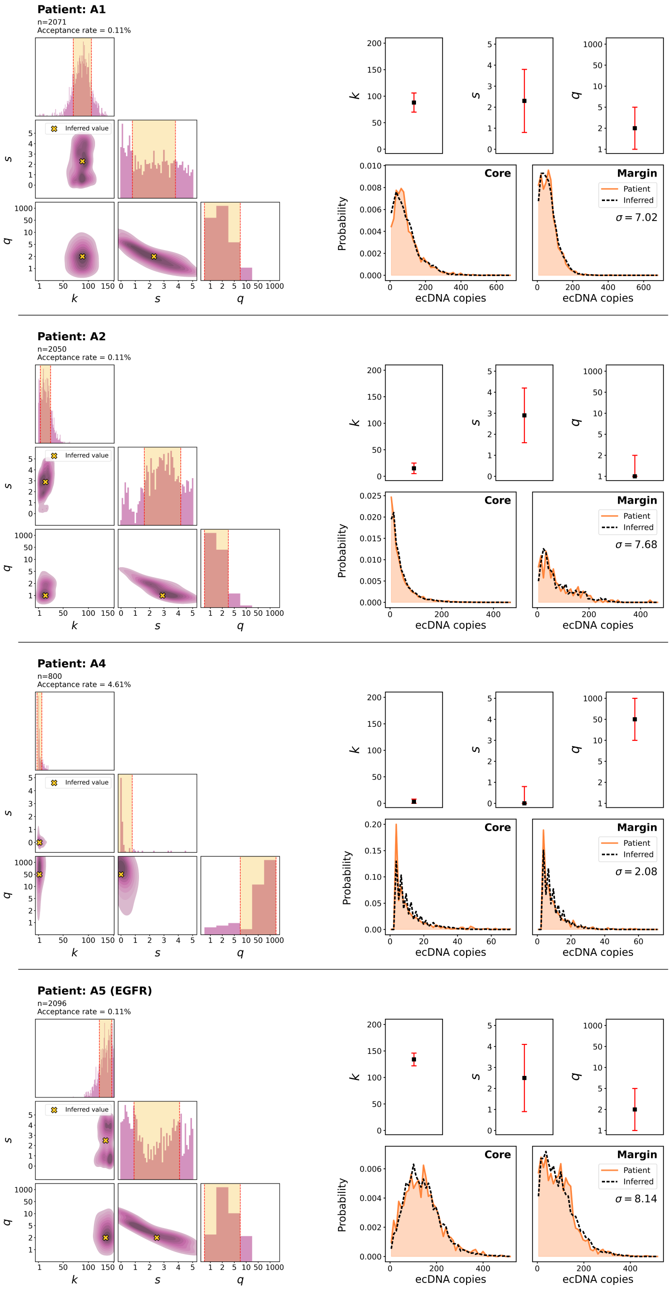


**Supplementary Figure 6:** *(Left)* Posterior parameter distributions for patients A1 to A5 (*EGFR*). Diagonal panels show 1D marginal distributions for $k$, $s$ and $q$. Off-diagonal show 2D marginal distributions for each combination of parameters. *(Right)* Summary of inferred $k$, $s$ & $q$ (top row) and patient-derived single-cell ecDNA copy number distribution, determined using DNA FISH, with corresponding best-fit distributions from simulated tumours (bottom row). Sum of Wasserstein distance between patient and simulated distributions for tumour core and infiltrating margin, representing closeness of fit, is denoted by $\sigma$.

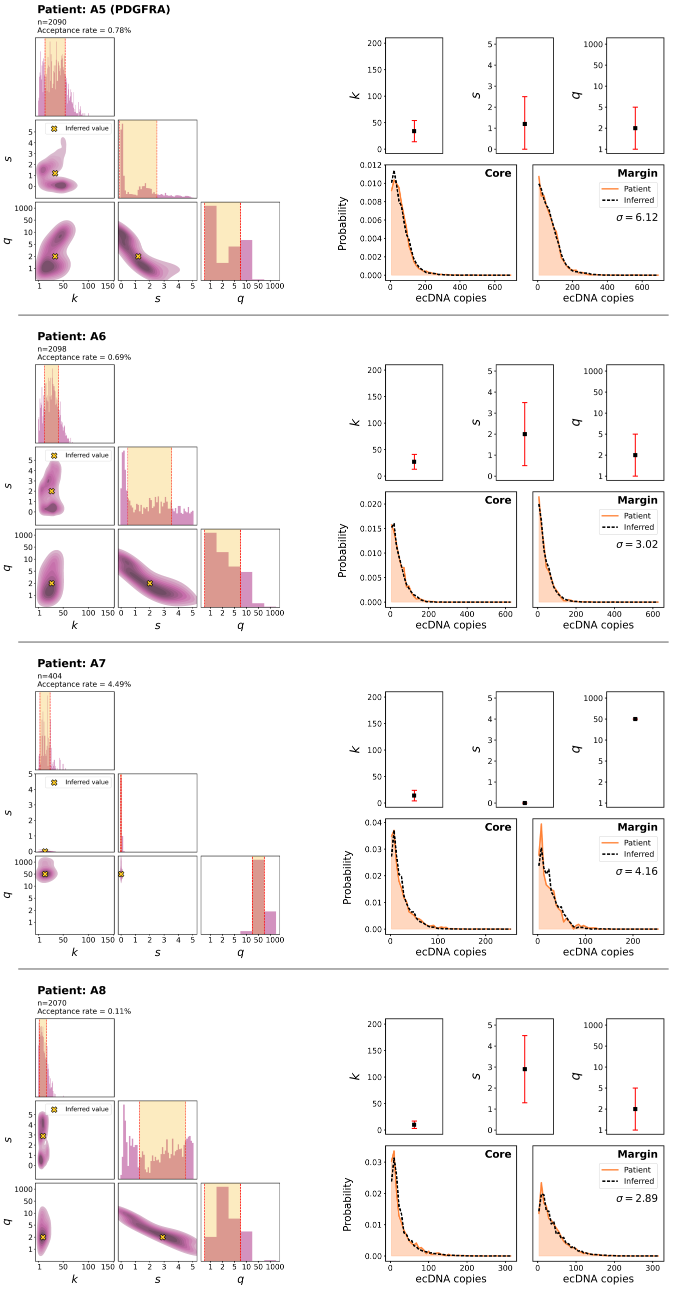


**Supplementary Figure 7:** *(Left)* Posterior parameter distributions for patients A5 (*PDGFRA*) to A8. Diagonal panels show 1D marginal distributions for $k$, $s$ and $q$. Off-diagonal show 2D marginal distributions for each combination of parameters. $n$ and acceptance rate denote the absolute and percentage number of simulations accepted into the posterior parameter set, respectively. *(Right)* Summary of inferred $k$, $s$ & $q$ (top row) and patient-derived single-cell ecDNA copy number distribution, determined using DNA FISH, with corresponding best-fit distributions from simulated tumours (bottom row). Sum of Wasserstein distance between patient and simulated distributions for tumour core and infiltrating margin, representing closeness of fit, is denoted by $\sigma$.

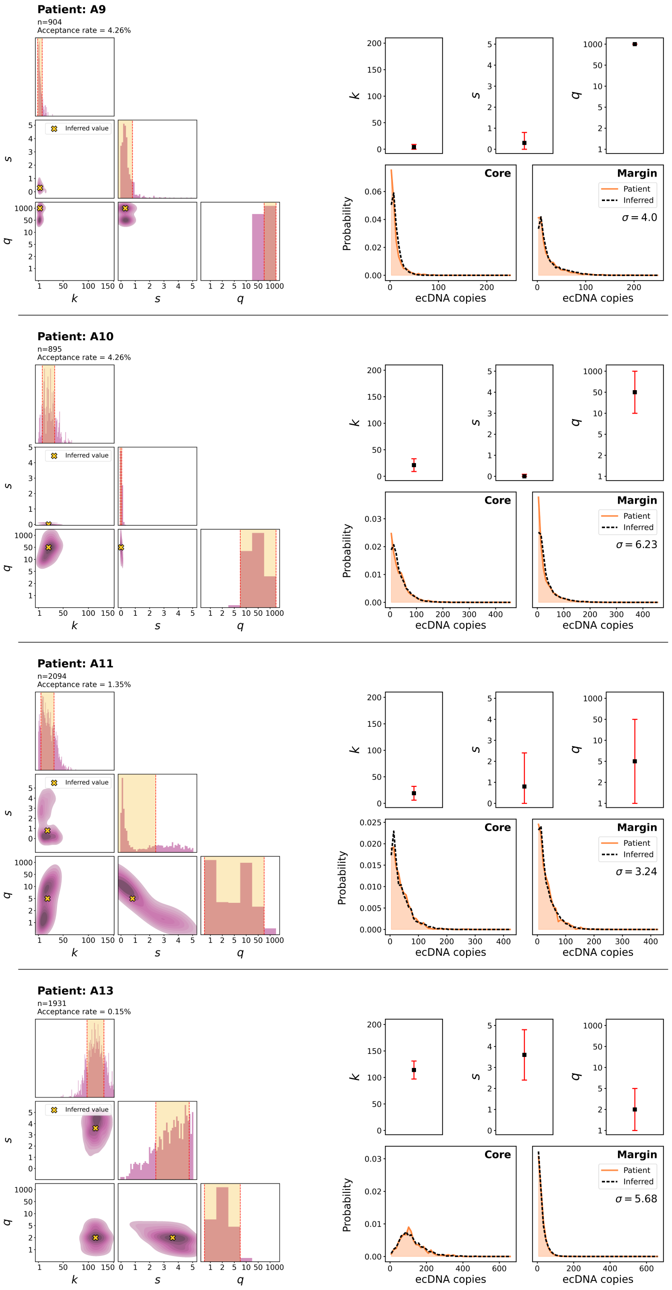


**Supplementary Figure 8:** *(Left)* Posterior parameter distributions for patients A9 to A13. Diagonal panels show 1D marginal distributions for $k$, $s$ and $q$. Off-diagonal show 2D marginal distributions for each combination of parameters. $n$ and acceptance rate denote the absolute and percentage number of simulations accepted into the posterior parameter set, respectively. *(Right)* Summary of inferred $k$, $s$ & $q$ (top row) and patient-derived single-cell ecDNA copy number distribution, determined using DNA FISH, with corresponding best-fit distributions from simulated tumours (bottom row). Sum of Wasserstein distance between patient and simulated distributions for tumour core and infiltrating margin, representing closeness of fit, is denoted by $\sigma$.

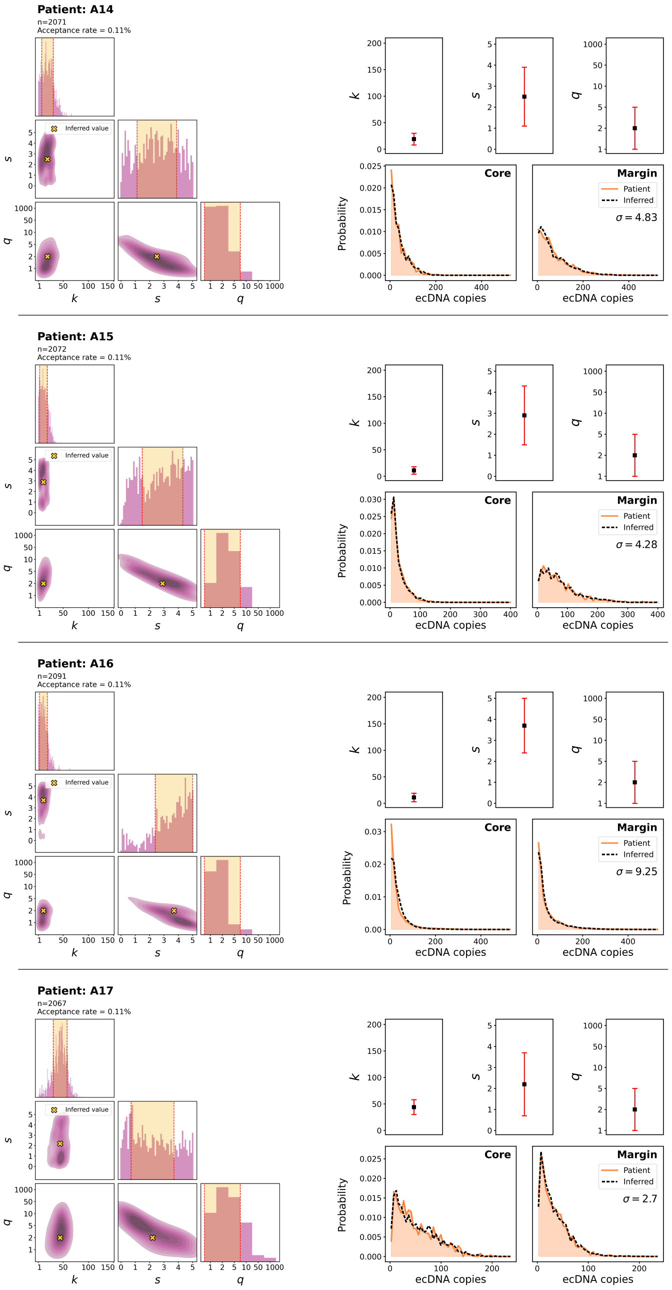


**Supplementary Figure 9:** *(Left)* Posterior parameter distributions for patients A14 to A17. Diagonal panels show 1D marginal distributions for $k$, $s$ and $q$. Off-diagonal show 2D marginal distributions for each combination of parameters. $n$ and acceptance rate denote the absolute and percentage number of simulations accepted into the posterior parameter set, respectively. *(Right)* Summary of inferred $k$, $s$ & $q$ (top row) and patient-derived single-cell ecDNA copy number distribution, determined using DNA FISH, with corresponding best-fit distributions from simulated tumours (bottom row). Sum of Wasserstein distance between patient and simulated distributions for tumour core and infiltrating margin, representing closeness of fit, is denoted by $\sigma$.

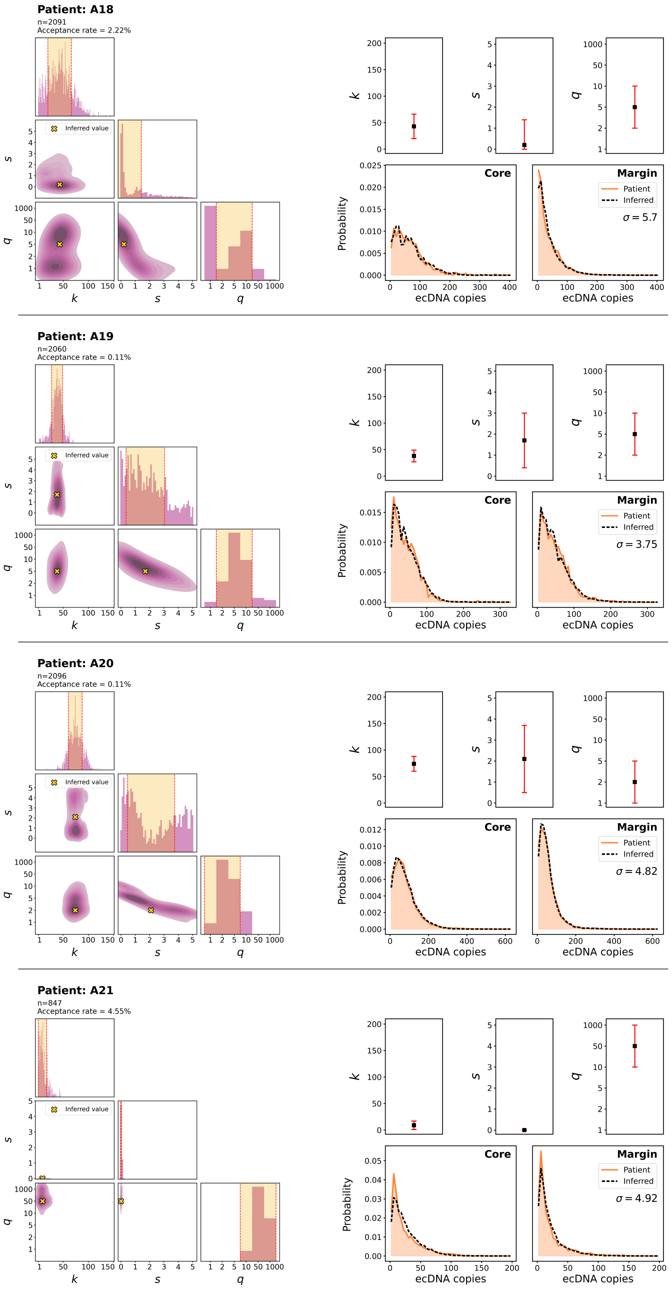


**Supplementary Figure 10:** *(Left)* Posterior parameter distributions for patients A18 to A21. Diagonal panels show 1D marginal distributions for $k$, $s$ and $q$. Off-diagonal show 2D marginal distributions for each combination of parameters. $n$ and acceptance rate denote the absolute and percentage number of simulations accepted into the posterior parameter set, respectively. *(Right)* Summary of inferred $k$, $s$ & $q$ (top row) and patient-derived single-cell ecDNA copy number distribution, determined using DNA FISH, with corresponding best-fit distributions from simulated tumours (bottom row). Sum of Wasserstein distance between patient and simulated distributions for tumour core and infiltrating margin, representing closeness of fit, is denoted by $\sigma$.

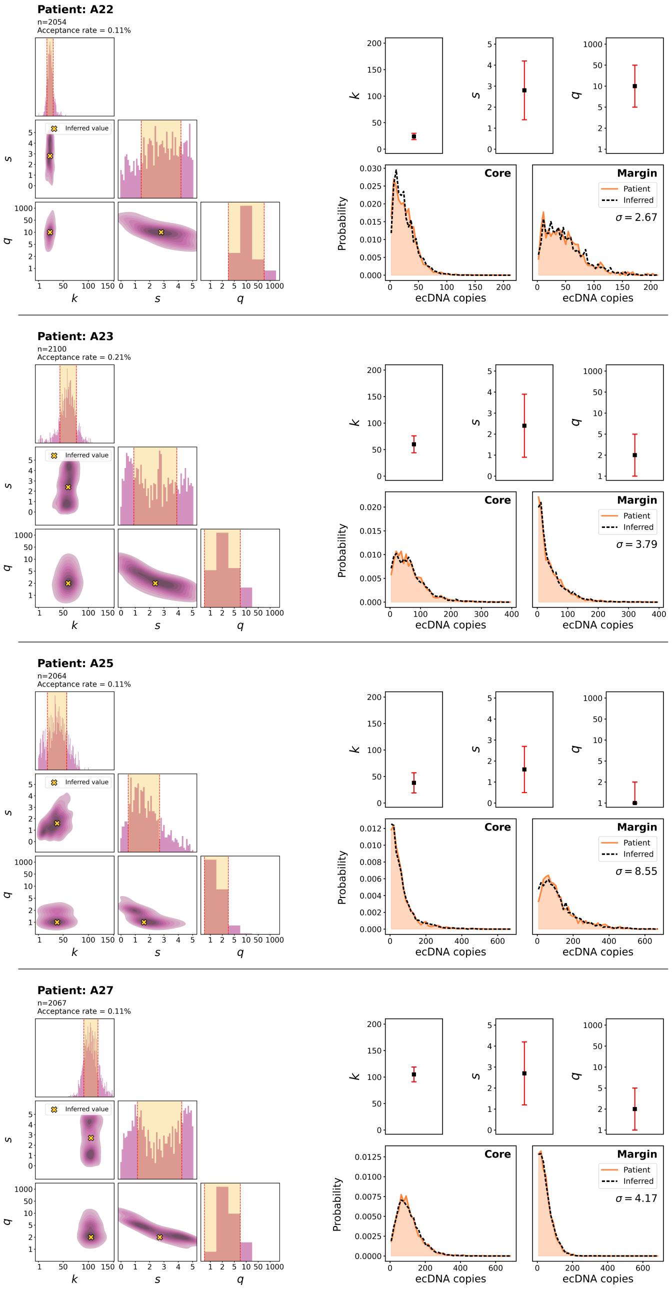


**Supplementary Figure 11:** *(Left)* Posterior parameter distributions for patients A22 to A27. Diagonal panels show 1D marginal distributions for $k$, $s$ and $q$. Off-diagonal show 2D marginal distributions for each combination of parameters. $n$ and acceptance rate denote the absolute and percentage number of simulations accepted into the posterior parameter set, respectively. *(Right)* Summary of inferred $k$, $s$ & $q$ (top row) and patient-derived single-cell ecDNA copy number distribution, determined using DNA FISH, with corresponding best-fit distributions from simulated tumours (bottom row). Sum of Wasserstein distance between patient and simulated distributions for tumour core and infiltrating margin, representing closeness of fit, is denoted by $\sigma$.

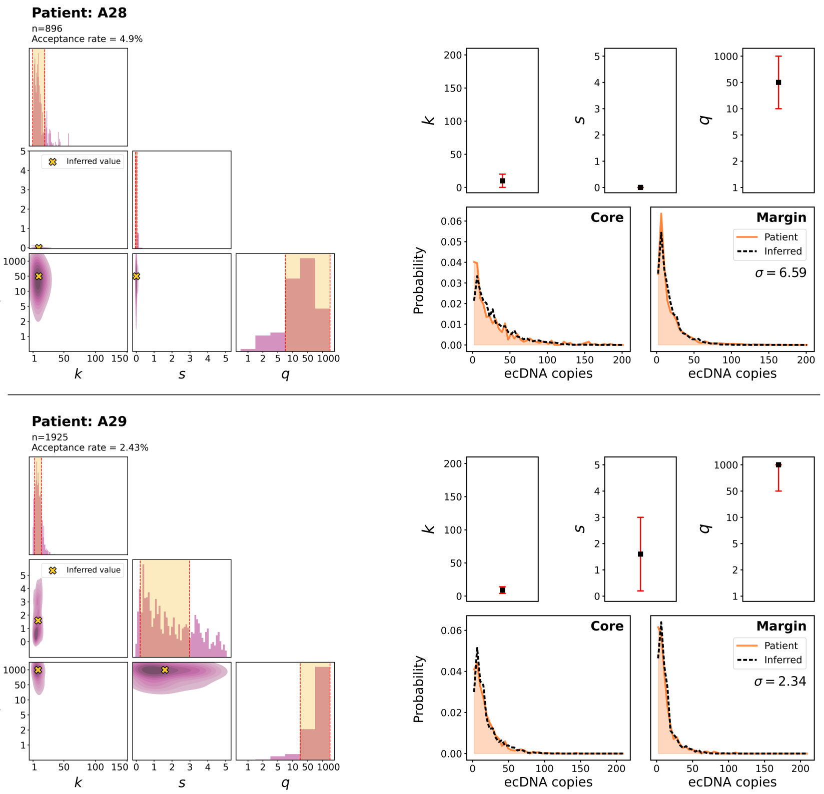


**Supplementary Figure 12:** *(Left)* Posterior parameter distributions for patients A28 and A29. Diagonal panels show 1D marginal distributions for $k$, $s$ and $q$. Off-diagonal show 2D marginal distributions for each combination of parameters. $n$ and acceptance rate denote the absolute and percentage number of simulations accepted into the posterior parameter set, respectively. *(Right)* Summary of inferred $k$, $s$ & $q$ (top row) and patient-derived single-cell ecDNA copy number distribution, determined using DNA FISH, with corresponding best-fit distributions from simulated tumours (bottom row). Sum of Wasserstein distance between patient and simulated distributions for tumour core and infiltrating margin, representing closeness of fit, is denoted by $\sigma$.

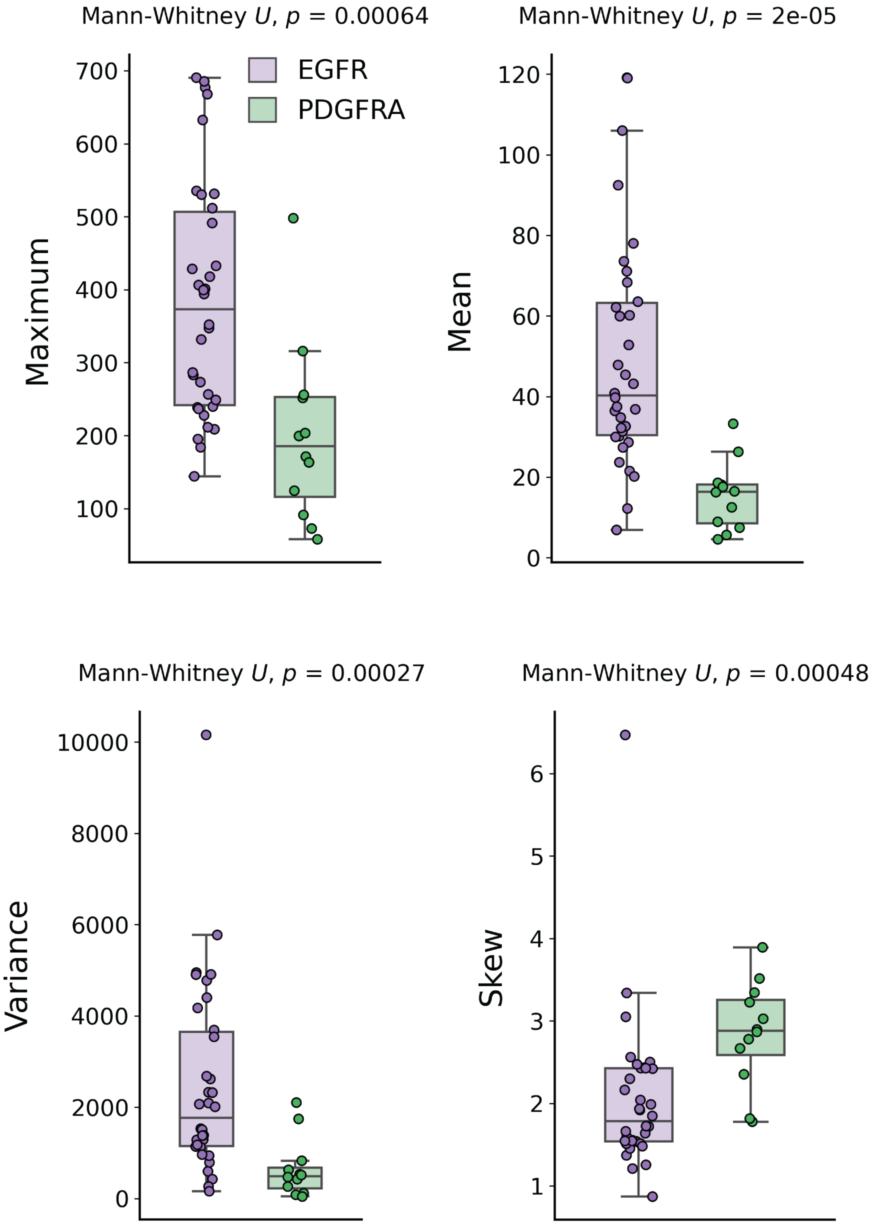


**Supplementary Figure 13:** Oncogene-level differences in observed ecDNA copy number distributions.

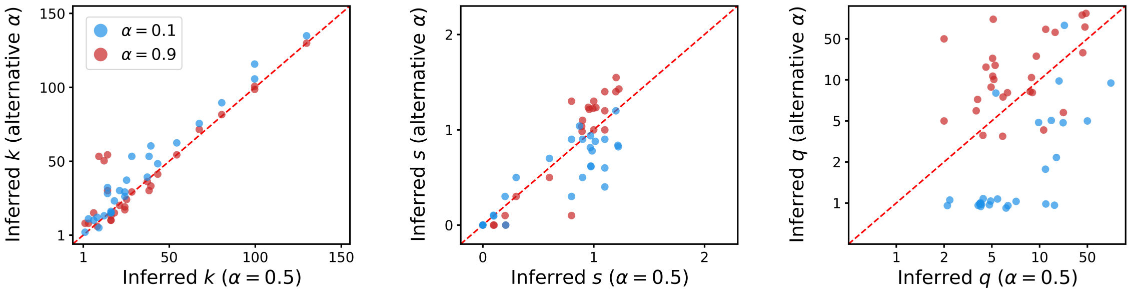


**Supplementary Figure 14:** Comparison of predicted model parameters when using a cell death rate of $\alpha = 0.5$ to alternative models using low ($\alpha = 0.1$) and high ($\alpha = 0.9$) death rates. Cell death rate, $r_d$, is computed as $r_d = r_b(x, 0) \cdot \alpha$, where $r_b(x, s = 0)$ denotes the neutral birth rate of a cell with $x$ copies of ecDNA.

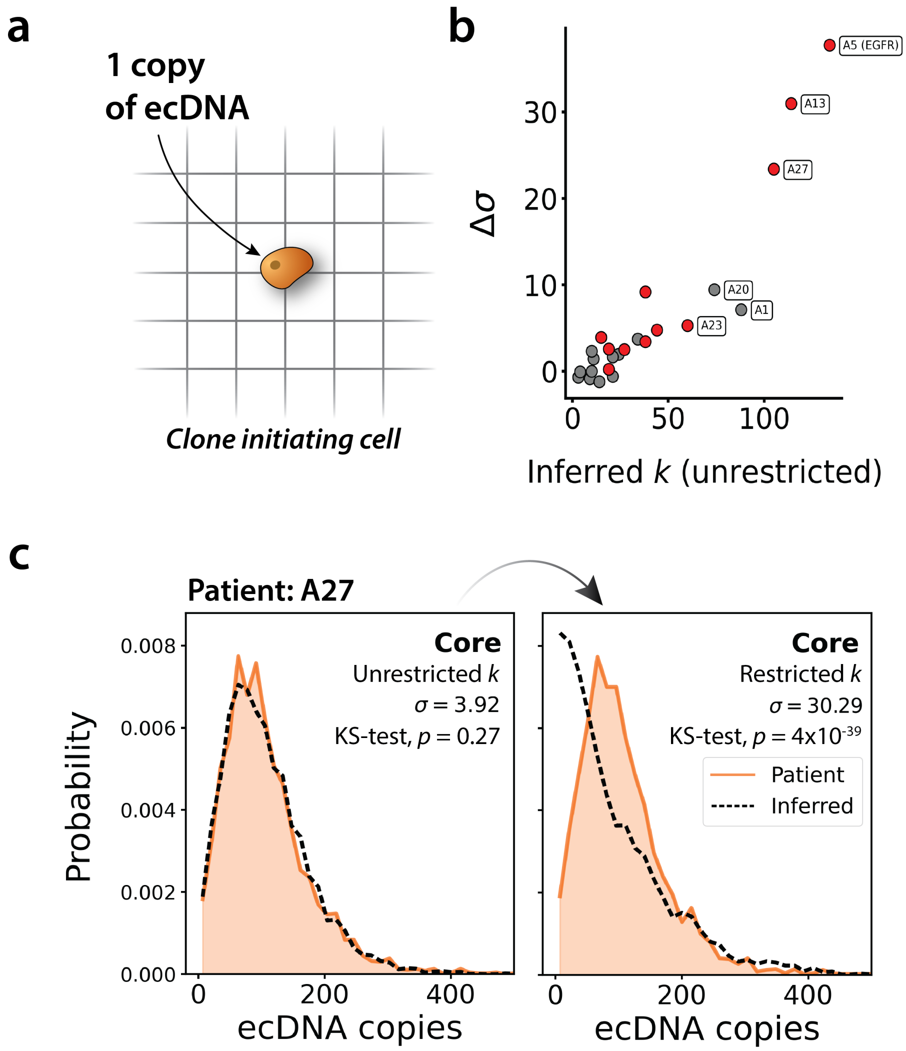


**Supplementary Figure 15:** **(a)** Parameter inference was repeated with a modified computational model, in which the initial tumour cell ecDNA copy number was restricted to $k = 1$. **(b)** ABC fit quality difference, $\Delta\sigma$, which compares inference with unrestricted $k$ value (Bayesian prior $P(k) = \mathrm{U}(1{,}150)$) to restricted case of $k = 1$. Larger values of $\Delta\sigma$ indicate a worse fit when $k$ is restricted. Red coloured points denote patients for which the best-fit model data was an exact fit to the patient data in the unrestricted $k$ case (core sample, Kolmogorov-Smirnov (KS) test, $p > 0.05$), but not in the restricted $k$ case (core sample, KS test, $p \leq 0.05$). **(c)** Example of poor model best-fit for patient A27 when $k$ is restricted to $k = 1$.

| | Sample | chromosome | position 1 | position 2 | sv_type | read support | amplicon type | spanning exons | vIII equivalent | c-terminal deletion |
|---|---|---|---|---|---|---|---|---|---|---|
| GB-UK | A5 | chr7 | 55200682 | 55205057 | deletion-like | 58 | ecDNA | 25-27 | | Y |
| | A8 | chr7 | 54920809 | 55154753 | inversion | 28 | ecDNA | 1-7 | Y | |
| | A11 | chr7 | 55132062 | 55155691 | deletion-like | 88 | ecDNA | 2-7 | Y | |
| | A14 | chr7 | 55119613 | 55154385 | inversion | 80 | ecDNA | 2-7 | Y | |
| | A15 | chr7 | 55118911 | 55155705 | deletion-like | 66 | ecDNA | 2-7 | Y | |
| | A16 | chr7 | 55021945 | 55179866 | inversion | 19 | ecDNA | 1-19 | | |
| | | chr7 | 55009553 | 55154336 | inversion | 23 | ecDNA | 1-7 | Y | |
| | A17 | chr7 | 55200774 | 55204714 | deletion-like | 38 | ecDNA | 25-27 | | Y |
| | A19 | chr7 | 55142134 | 55154633 | deletion-like | 36 | ecDNA | 2-7 | Y | |
| | | chr7 | 54843745 | 55128844 | inversion | 20 | ecDNA | 1 | | |
| | | chr7 | 54880847 | 55154597 | inversion | 21 | ecDNA | 1-7 | Y | |
| | | chr7 | 54884387 | 55141732 | inversion | 17 | ecDNA | 1 | | |
| | A20 | chr7 | 55025248 | 55155160 | deletion-like | 4 | ecDNA | 2-7 | Y | |
| | | chr7 | 55122733 | 55154336 | deletion-like | 4 | ecDNA | 2-7 | Y | |
| | | chr7 | 55123695 | 55155612 | deletion-like | 6 | ecDNA | 2-7 | Y | |
| | A23 | chr7 | 55026802 | 55155149 | deletion-like | 5 | ecDNA | 2-7 | Y | |
| | | chr7 | 55132447 | 55153405 | foldback | 2 | ecDNA | 2-6 | | |
| | A27 | chr7 | 55201858 | 56259403 | inversion | 7 | ecDNA | 27,28+ | | Y |
| | | chr7 | 55202026 | 55742391 | deletion-like | 10 | ecDNA | 27,28+ | | Y |
| | A29 | chr7 | 55093609 | 55155808 | deletion-like | 17 | ecDNA | 2-7 | Y | |
| PCAWG | DO12952-SP27603 | chr7 | 55097880 | 55223306 | deletion-like | 54 | ecDNA | 2-7 | Y | |
| | DO11238-SP24301 | chr7 | 55199311 | 55222950 | deletion-like | 45 | ecDNA | 2-7 | Y | |
| | | chr7 | 55199544 | 55222330 | deletion-like | 309 | ecDNA | 2-7 | Y | |
| | DO10960-SP23739 | chr7 | 55122639 | 55223386 | deletion-like | 24 | ecDNA | 2-7 | Y | |
| | | chr7 | 55130764 | 55223089 | deletion-like | 24 | ecDNA | 2-7 | Y | |
| | | chr7 | 55268961 | 55274292 | deletion-like | 15 | ecDNA | 25-28 | | Y |
| | DO12034-SP25833 | chr7 | 55105714 | 55221814 | deletion-like | 410 | ecDNA | 2-7 | Y | |
| | DO13192-SP28041 | chr7 | 55191873 | 55223395 | deletion-like | 86 | ecDNA | 2-7 | Y | |
| | | chr7 | 55201724 | 55222811 | deletion-like | 131 | ecDNA | 2-7 | Y | |
| | DO12454-SP26649 | chr7 | 54958584 | 55223150 | inversion | 106 | ecDNA | 2-7 | | |
| | DO11202-SP24236 | chr7 | 55150598 | 55221850 | deletion-like | 53 | ecDNA | 2-7 | Y | |
| | DO13474-SP28581 | chr7 | 55198927 | 55222711 | deletion-like | 4 | ecDNA | 2-7 | Y | |
| | DO11854-SP25494 | chr7 | 55146310 | 55222238 | deletion-like | 57 | ecDNA | 2-7 | Y | |

**Supplementary Figure 16:** Structural variant analysis of EGFR ecDNA in GB-UK and PCAWG GBM samples.

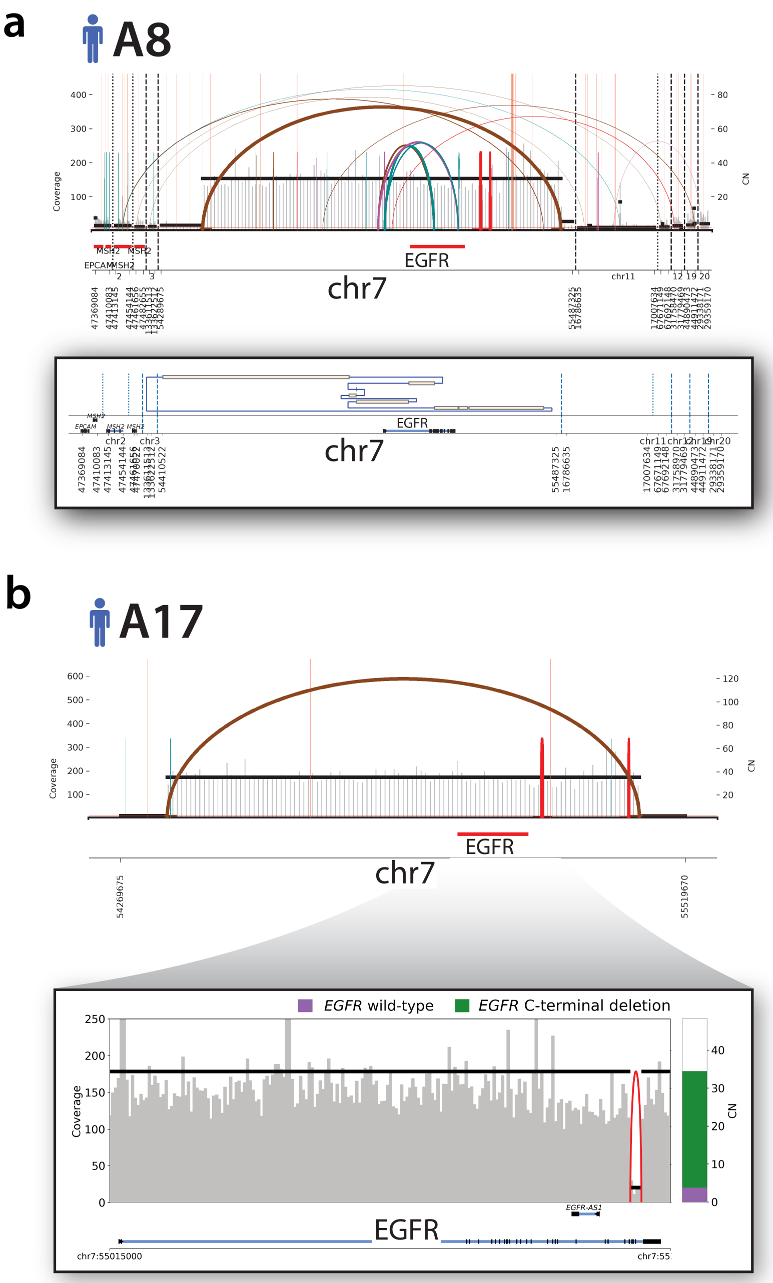


**Supplementary Figure 17:** Representative structural variants in *EGFR*-ecDNA. **(a)** Upper panel is a Sashimi plot highlighting the structural variants and relative copy number of *EGFR*-ecDNA in patient A8. Lower panel is a joining plot showing a translocation of EGFR exons 1 – 7 to upstream of *EGFR* in an inverted orientation. **(b)** Upper panel is a Sashimi plot highlighting a simple circular *EGFR*-ecDNA in patient A17. Lower panel is a zoomed in view of *EGFR* showing a deep c-terminal deletion within the ecDNA.

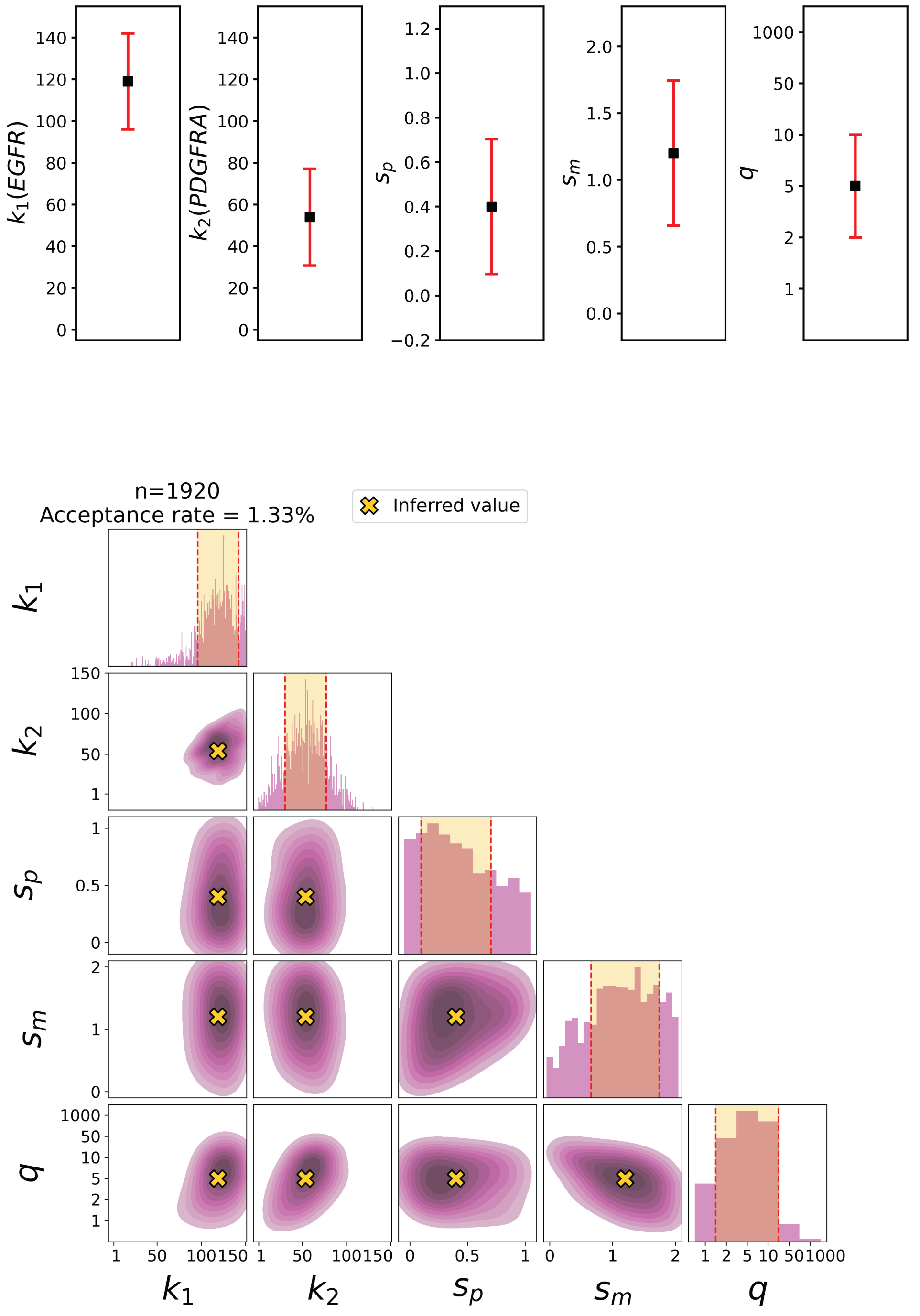


**Supplementary Figure 18:** Model inference summary for patient A5 using the co-amplified ecDNA simulation model. Model was fit to ecDNA copy number distributions for *EGFR*-amplifying and *PDGFRA*-amplifying ecDNA species. Parameters $k_1$ and $k_2$ describe the initial number of each ecDNA species, whilst $s_p$ and $s_m$ represent selection coefficients for tumour cells with 1 (pure) or 2 (mixed) ecDNA species respectively.

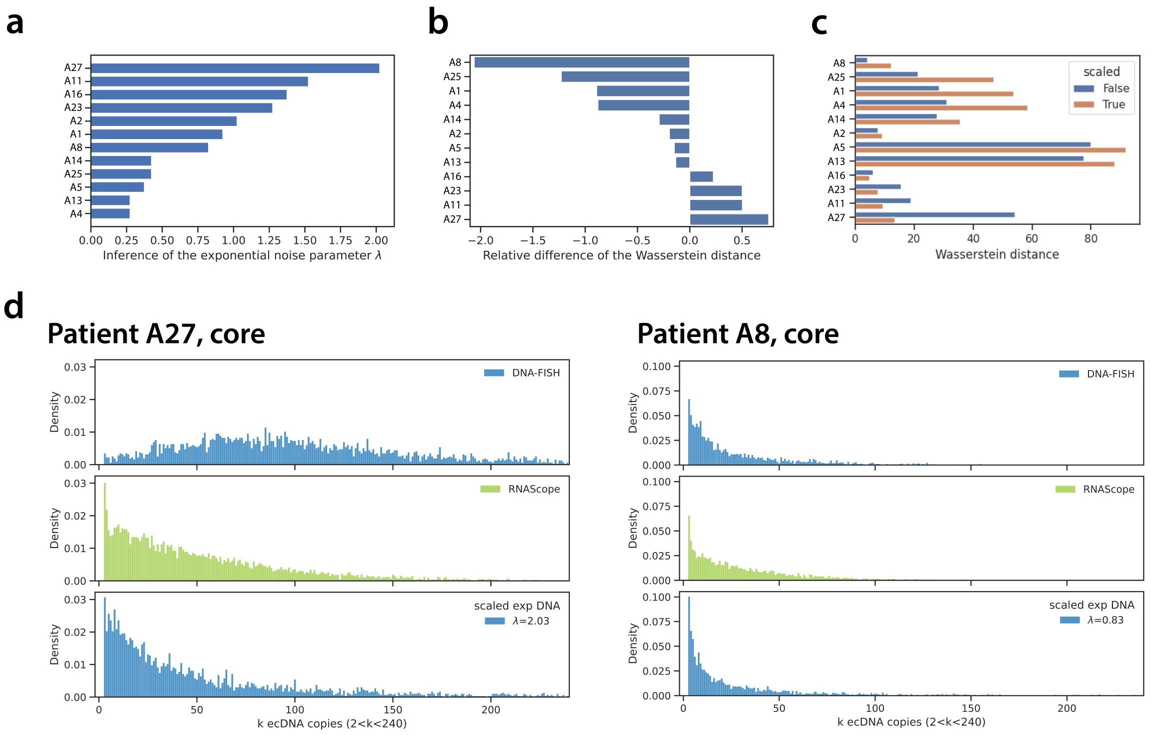


**Supplementary Figure 19:** Concordance of per-cell ecDNA distributions, derived with DNA FISH, and nascent RNAscope distributions for a range of patient samples. **(a)** Parameter of exponential noise kernel, $\lambda$, which provided best fitting transformation of DNA to RNA distributions. Best-fitting values were inferred using approximate Bayesian computation. **(b)** Difference between re-scaled DNA and RNA distributions, relative to distance between original DNA and RNA distributions. **(c)** Wasserstein distance between DNA and RNA distributions, before and after scaling of the DNA distribution. **(d)** Examples for two patients, A27 and A8, of the experimentally measured DNA and RNA distributions (top and middle row respectively) and the re-scaled DNA distribution (bottom).

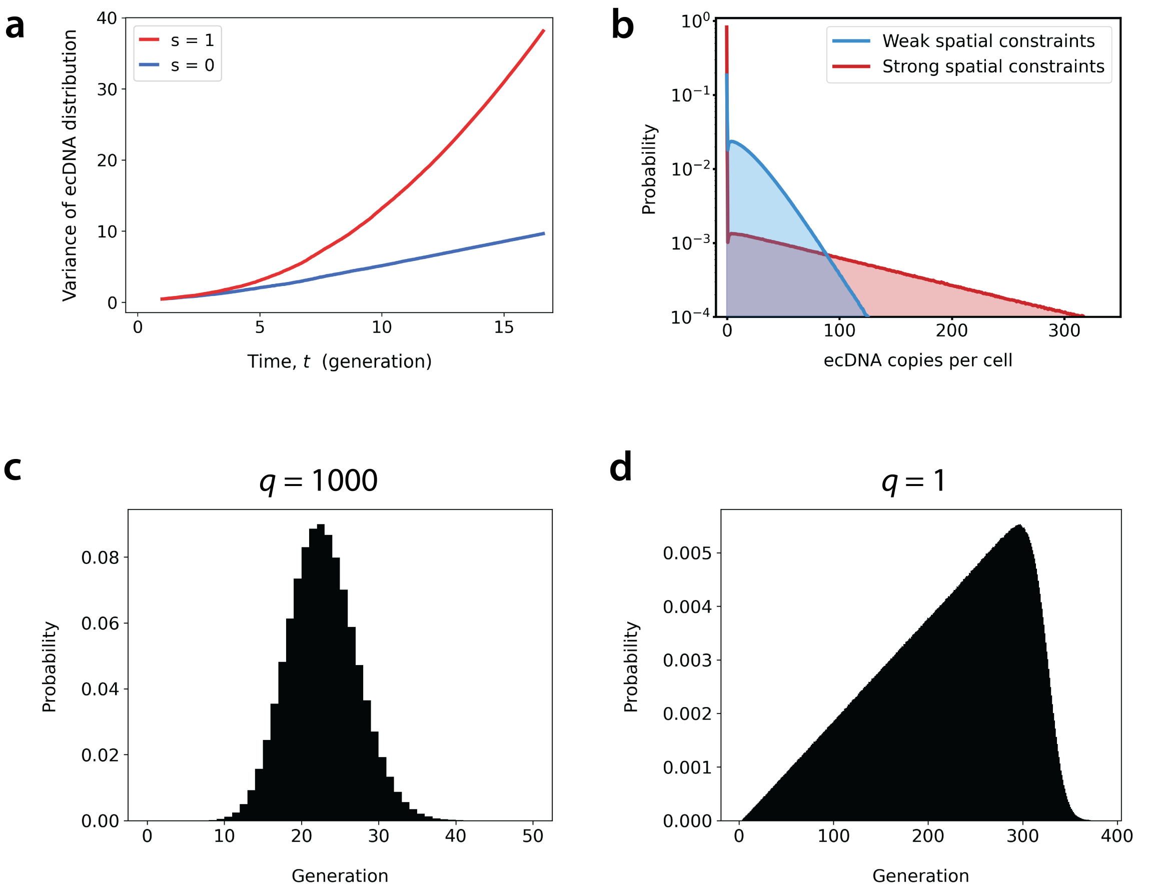


**Supplementary Figure 20:** **(a)** Variance in single-cell ecDNA copy number as a function of time for non-spatially constrained tumours ($q = 1000$) with initial ecDNA copy number $k = 1$. Variance scales linearly with time when ecDNA presence confers no fitness advantage ($s = 0$) and quadratically when ecDNA give rise to increased cell fitness ($s = 1$). **(b)** EcDNA copy number distribution for tumours with weak ($q = 1000$) versus strong ($q = 5$) spatial constraints ($k = 20$ & $s = 0$). Distributions represent combined data from 500 simulated tumours, final tumour size $N_{max} = 10^5$ cells. **(c-d)** Cell generation (number of previous cell divisions in cell lineage) distributions for tumours with **(c)** $q = 1000$ and **(d)** $q = 1$, simulated up to final a size $N_{max} = 10^5$ cells. Distributions represent combined data from 500 simulated tumours.

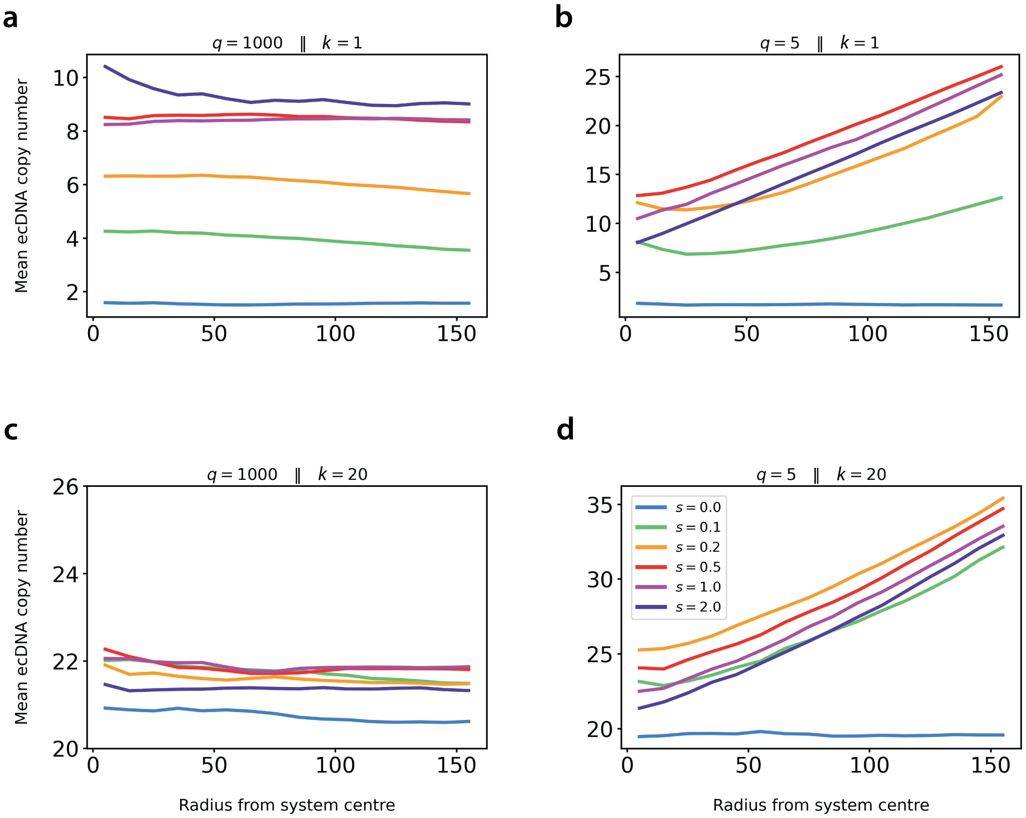


**Supplementary Figure 21:** Relationship between mean single-cell ecDNA copy number with distance from tumour centre over a range of ecDNA fitness advantages, $s$ for tumours with **(a)** $q = 1000$ & $k = 1$; **(b)** $q = 5$ & $k = 1$; **(c)** $q = 1000$ & $k = 20$; **(d)** $q = 5$ & $k = 20$. Data represent averaged values from 500 simulated tumours, final tumour size $N_{max} = 10^5$ cells.

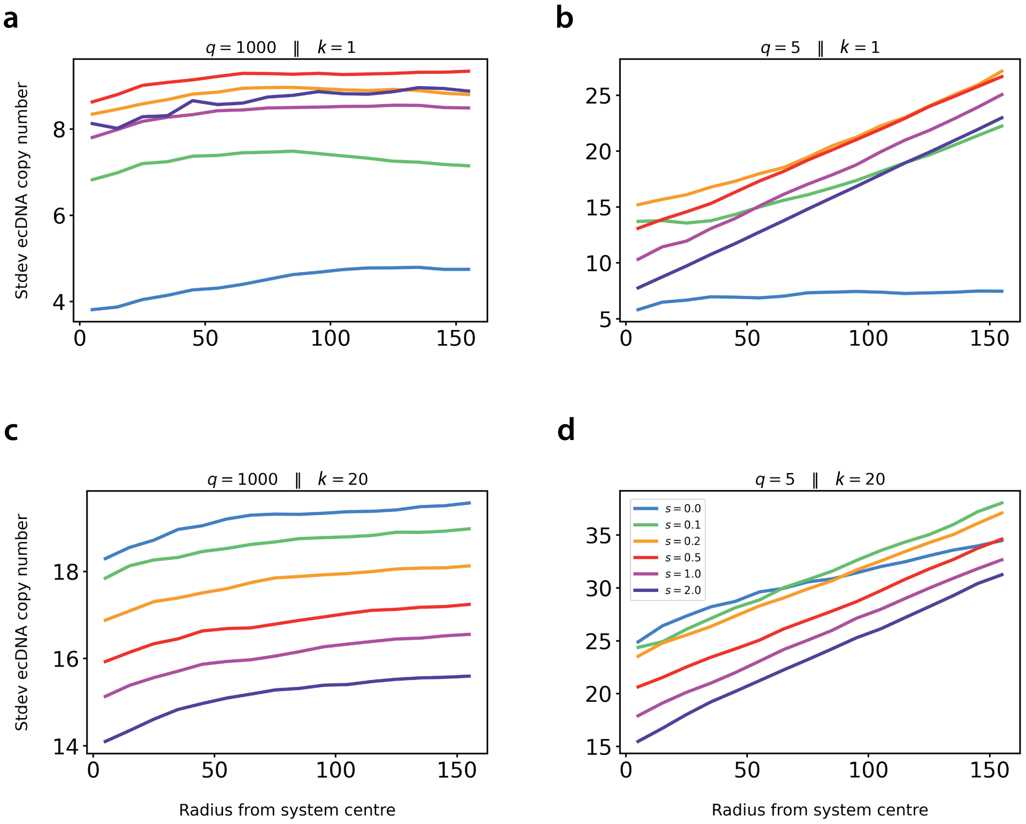


**Supplementary Figure 22:** Relationship between the standard deviation in single-cell ecDNA copy number with distance from tumour centre over a range of ecDNA fitness advantages, $s$ for tumours with **(a)** $q = 1000$ & $k = 1$; **(b)** $q = 5$ & $k = 1$; **(c)** $q = 1000$ & $k = 20$; **(d)** $q = 5$ & $k = 20$. Data represent averaged values from 500 simulated tumours, final tumour size $N_{max} = 10^5$ cells.

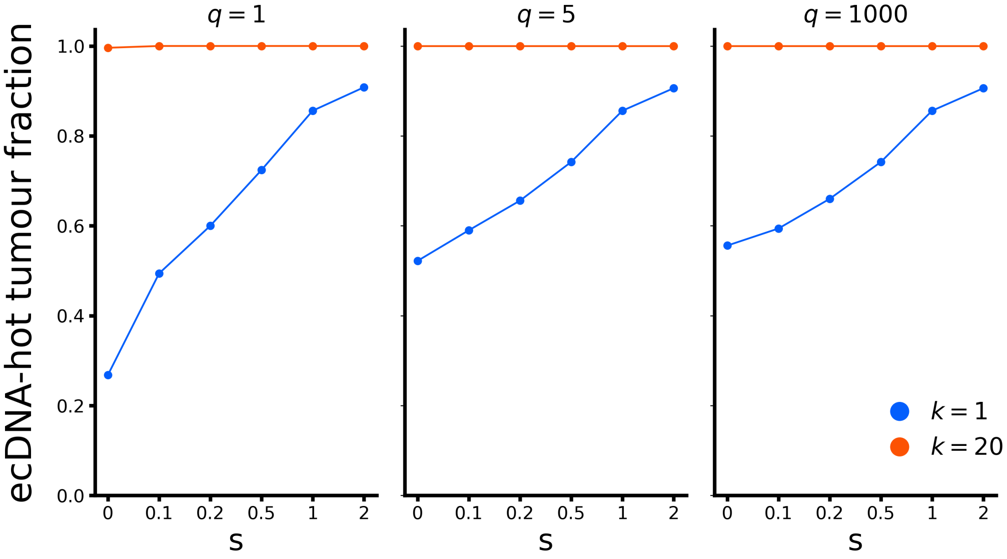


**Supplementary Figure 23:** Fraction of simulated tumours which maintain ecDNA within at least one tumour cell (termed ecDNA-hot tumours) measured at a final tumour size of $N_{max} = 10^5$ cells. Data represent mean values for 500 independent realisations of the stochastic simulation.

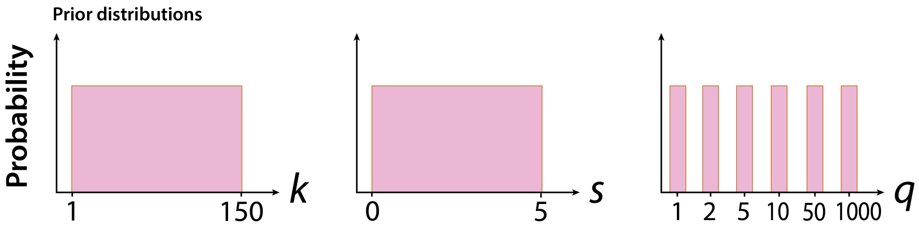


**Supplementary Figure 24:** Prior distributions for model parameters $k$, $s$ and $q$, used for approximate Bayesian computation.

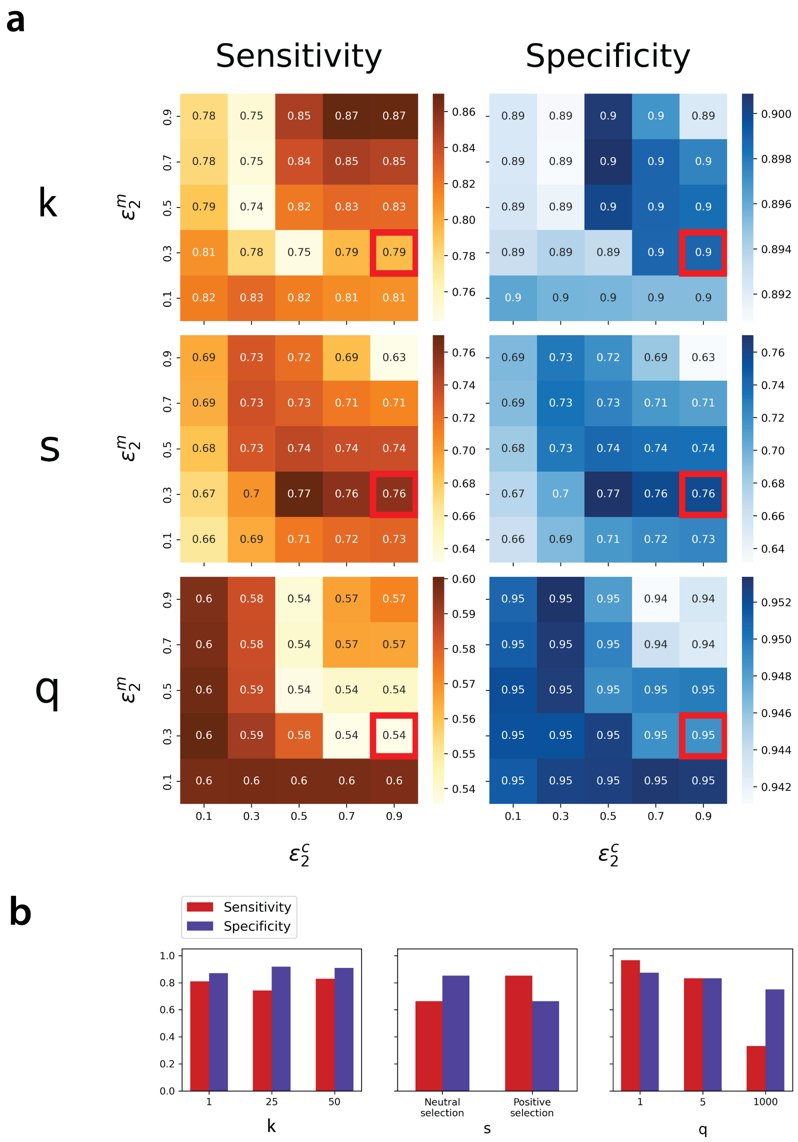


**Supplementary Figure 25:** **(a)**Averaged sensitivity and specificity of model inference algorithm using a range of threshold values on low-ecDNA cell fraction similarity, $\epsilon_2^c$ and $\epsilon_2^m$, for the tumour core and margin respectively. Red boxes highlight combination of threshold values which maximised algorithm sensitivity and specificity. **(b)** Sensitivity (true positive rate) and specificity (true negative rate) of ABC inference algorithm tested across a range of model $k$, $s$ and $q$ values. The algorithm was considered to have predicted a given value if that value was contained within the interval of the point estimator $\pm$ error. Data were generated by running the ABC algorithm on artificial patient core and margin ecDNA data, generated themselves using the spatial model. Artificial patient dataset represented all possible combinations of $k \in \{1, 25, 50, 100\}$, $s \in \{0, 0.5, 1, 2\}$ and $q \in \{1, 5, 50, 1000\}$, with 100 patients per parameter combination.

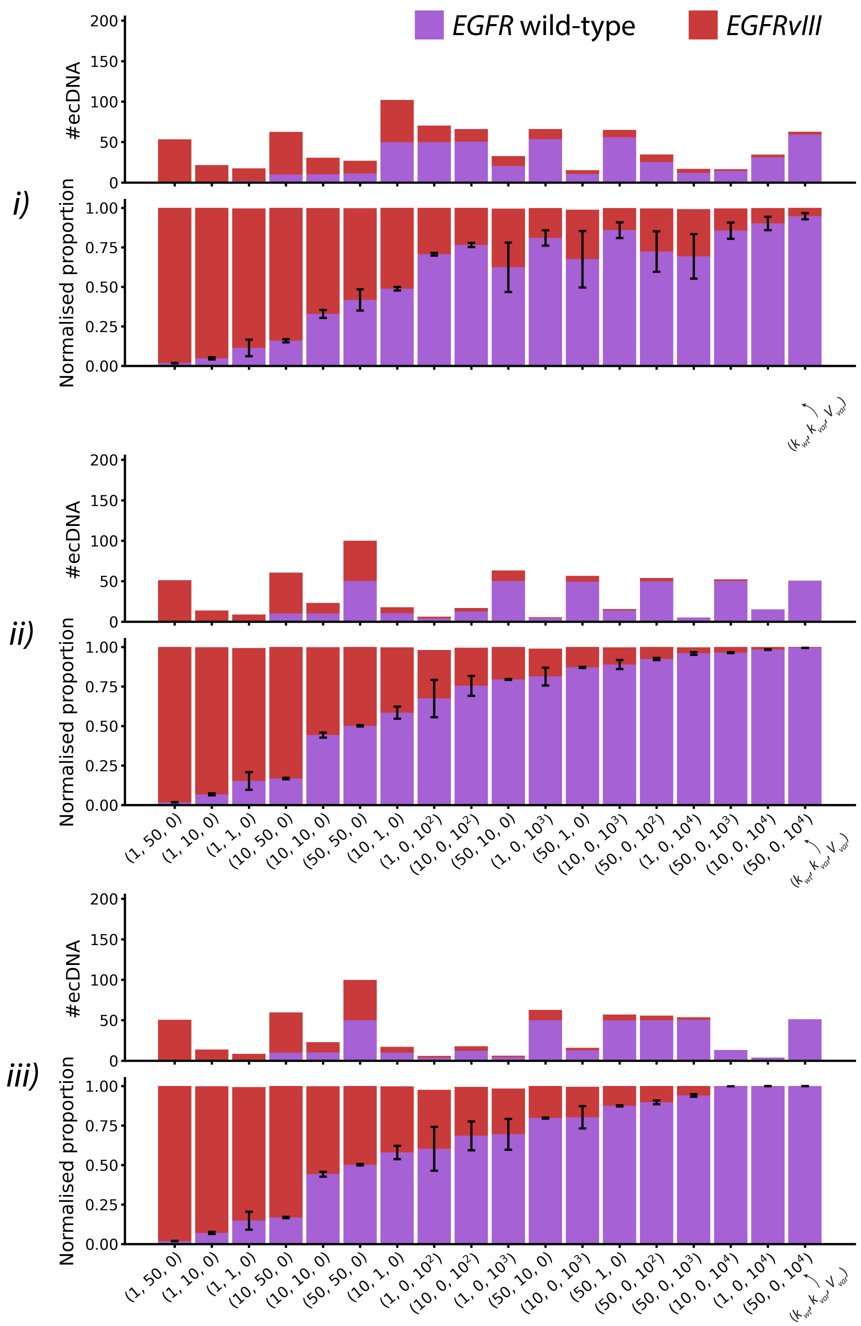


**Supplementary Figure 26:** Mean core ecDNA heteroplasmy in final tumour of $N_{max} = 10^5$ cells, simulated for a range of $(k_{wt}, k_{var}, V_{var})$ values with $s_{wt} = 0.2$, $s_{var} = 2$, and $(i)$ $q = 2$; $(ii)$ $q = 10$ or $(iii)$ $q = 1,000$. In each panel, top and bottom rows contain absolute and percentage numbers of ecDNA carrying wild-type or mutated *EGFR*. Values are mean ± standard deviation, derived from 1,000 simulated tumours.

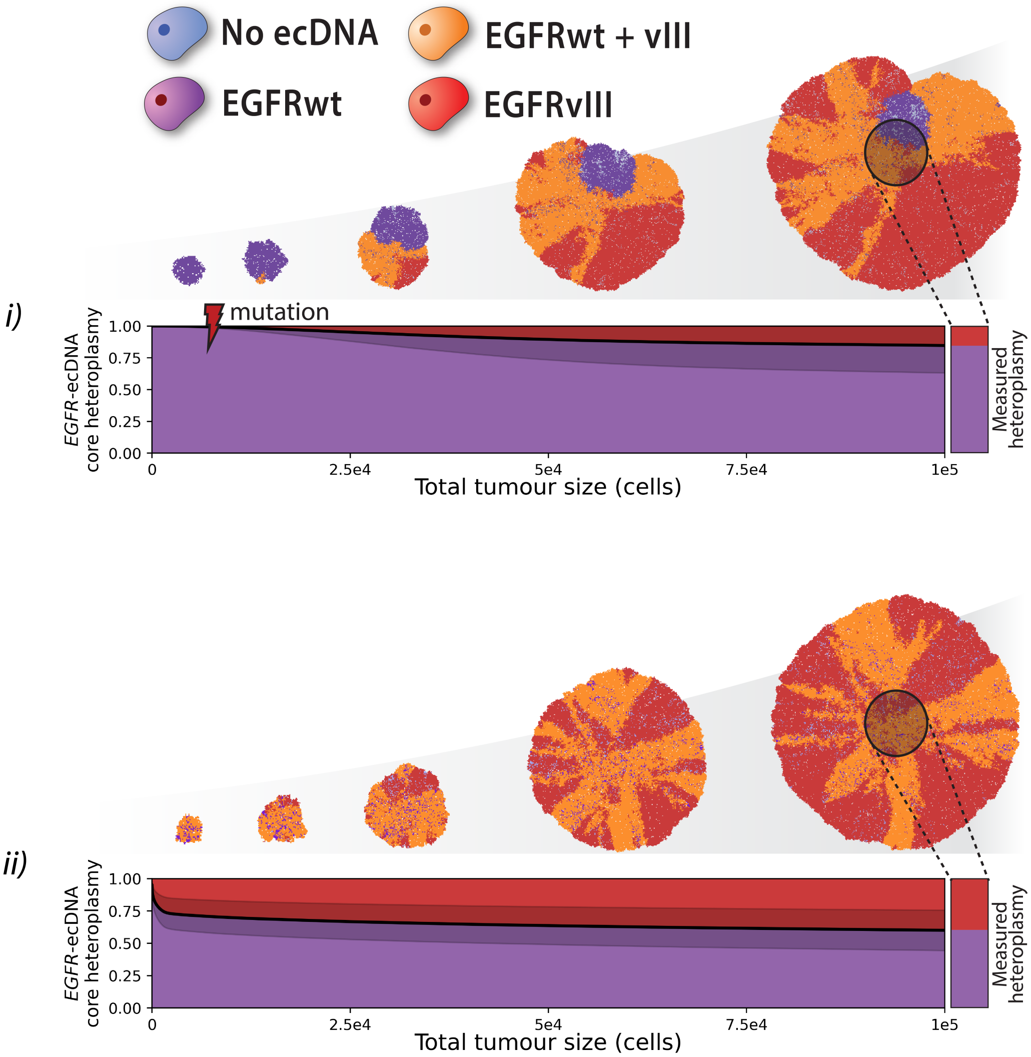


**Supplementary Figure 27:** Time-evolution of ecDNA heteroplasmy in tumour core for $(i)$ ($k_{wt} = 50$, $k_{var} = 0$, $V_{var} = 1,000$) and $(ii)$ ($k_{wt} = 20$, $k_{var} = 1$, $V_{var} = 0$). Final tumour size of $N_{max} = 10^5$ cells, and $s_{wt} = 0.2$, $s_{var} = 2$, $q = 2$. Upper row depicts time-evolution of single representative tumour; lower row represent mean ± standard deviation (shaded region) from 1,000 simulated tumours.

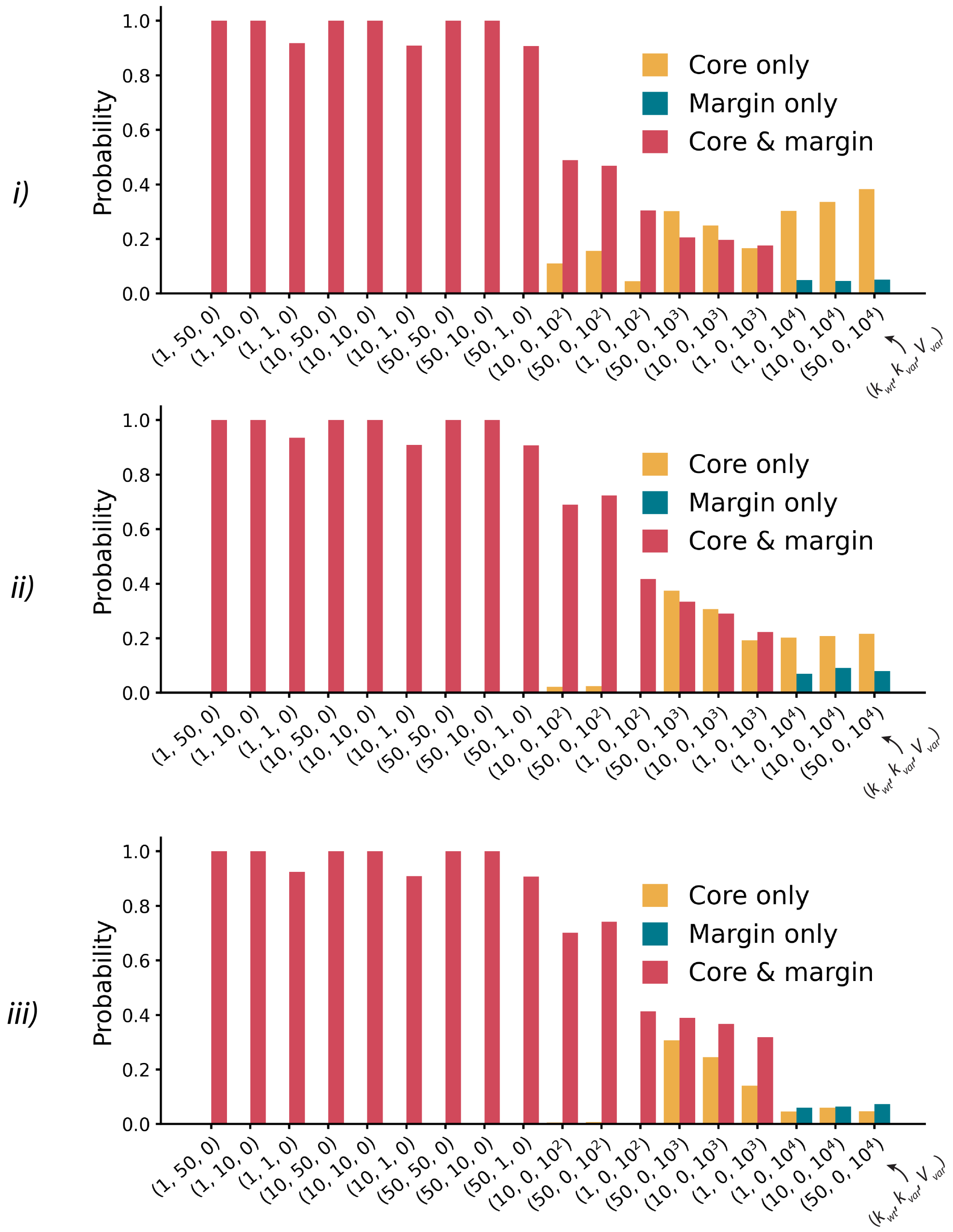


**Supplementary Figure 28:** Probability of observing variant ecDNA across tumour core and margin, for a pair of core and random margin regions in final tumour of $N_{max} = 10^5$ cells, simulated for a range of $(k_{wt}, k_{var}, V_{var})$ values with $s_{wt} = 0.2$, $s_{var} = 2$, and $(i)$ $q = 2$; $(ii)$ $q = 10$ or $(iii)$ $q = 1,000$.. Mean values derived from 1,000 simulated tumours.
